# Towards a probiotic approach for building plumbing – nutrient-based selection during initial biofilm formation on flexible polymeric materials

**DOI:** 10.1101/2020.04.10.033217

**Authors:** Neu Lisa, Cossu Laura, Frederik Hammes

**Author notes:** Corresponding author: **Name:** Frederik Hammes, **Email:**.

## Abstract

Upon entering building plumbing systems, drinking water bacteria experience considerable changes in environmental conditions. For example, some flexible polymeric materials leach organic carbon, which increases bacterial growth and reduces diversity. Here we show that the carbon supply by a flexible polymeric material drives nutrient-based selection within establishing biofilm communities. We found that migrating carbon from EPDM coupons resulted in considerable growth for different drinking water communities (0.2 – 3.3 × 10^8^ cells/cm^2^). All established biofilm communities showed low diversity (29 – 50 taxa/biofilm), with communities dominated by even viewer taxa (e.g., 5 taxa accounting for 94 ± 5 % relative abundance, n = 15). Interestingly, biofilm communities shared some taxa (e.g., *Methylobacterium* spp.) and families (e.g., Comamonadaceae), despite the difference in starting communities. Moreover, selected biofilm communities performed better than their original communities regarding maximum attachment (91 ± 5 vs. 69 ± 23 %, n = 15) and attachment rate (5.0 ± 1.7 × 10^4^ vs. 2.4 ± 1.2 × 10^4^ cells/cm^2^/h, n = 15) when exposed to new EPDM coupons. Our results demonstrate nutrient-based selection during initial biofilm formation on a flexible polymeric material and a resulting benefit to selected communities. We anticipate our findings to help connecting observational microbiological findings with their underlying ecological principles. Regarding initial biofilm formation, attachment dynamics, growth, and selection thereof are important for the management of microbial communities. In fact, managing initial colonization by supplying specific carbon and/or introducing consciously chosen/designed communities potentially paves the way for a probiotic approach for building plumbing materials.

## 1. Introduction

Uncontrolled microbial growth in building plumbing systems is generally undesirable as it can lead to operational and/or hygienic problems^1,2^. Such growth is caused by changes in environmental conditions, which is what drinking water bacteria experience as soon as they enter a building plumbing system. For example, water temperature increases and fluctuates spatially and temporally, which was shown to alter community composition^3,4^. Also, pipe diameters are considerably smaller (e.g., < 2 cm) compared to main distribution pipes (e.g., ≥ 10 cm), which provides more surface area per water volume^5^, and increases the impact of biofilms on the water phase. Regarding operation, flow pattern and rates have been shown to impact biofilm structure and community composition^6,7^. Finally, diverse materials are used for pipes and non-pipe components^8^, and some of these support microbial growth by leaching biodegradable substances^9^, which is especially critical under long stagnation times of the water^10^. The bottom line is that building plumbing systems often provide more favorable environmental conditions for bacterial growth than the main distribution network and that it is important to understand and control not only their individual but also their combined impact on the drinking water microbiome.

Several previous studies investigated the impact of building plumbing conditions on its microbiome. Overall, microbial community compositions tend to change considerably, e.g., (1) during stagnation^11^, (2) while forming biofilms inside flexible shower hoses^12^, or (3) due to the combined impact of material, temperature, and stagnation^13^. Considering one of the above in more detail, studies in our research group that were addressing biofilm formation on flexible polymeric materials revealed (1) high bacterial numbers (i.e., growth) and (2) a considerable loss in species diversity (i.e., selection)^12,14^. Also, Proctor and colleagues^15^ observed the development of dissimilar biofilm community compositions when exposing the same drinking water community to different polymeric hose materials. Thereupon, they reasoned for considerable impact of migrating organic carbon on both growth and selection.

In this study, we investigated nutrient-based selection during initial biofilm formation, using a microcosm set up for the simulation of new flexible polymeric material (EPDM) in contact with drinking water. Our hypotheses were: (1) EPDM coupons release biodegradable organic carbon, which increases the potential of bacteria to grow in an otherwise carbon-limited environment. (2) Selection occurs within establishing biofilm communities, irrespective of the initial drinking water community composition. (3) Due to the common carbon supply, biofilm communities will show a certain degree of similarity in their compositions. (4) The selection process will bring advantages for initial biofilm formation, e.g., attachment, growth, etc.. We finally argue that this information provides an opportunity for the development of new, pro-active approaches for the management of biofilms that form on polymeric building plumbing materials.

## 2. Materials and Methods

### 2.1 Selection of coupon material and water

The interplay between one flexible polymeric material and five different drinking waters was tested regarding biofilm formation and community selection (Figure 1A). Coupons of ethylene propylene diene monomer (EPDM) rubber (Angst+Pfister AG, Switzerland) with an ethylene fraction of 2 % (w/w) was used as experimental material throughout this study. EPDM is approved for the use in contact with drinking water^16,17^, e.g. as rubber seals within building plumbing systems. Two bottled waters and three non-chlorinated tap waters were selected as water matrices, namely: Evian (France; *B1*), Aproz (Switzerland; *B2*), tapped groundwater Dübendorf (Switzerland; *T1*), tap water Dübendorf (Switzerland; *T2*), and tap water Oerlikon (Switzerland; *T3*). The five waters differed in their chemical and biological composition. However, all waters were oligotrophic with low organic carbon and phosphorous concentrations and total bacterial concentrations in the same order of magnitude (1 – 3 × 10^5^ cells/mL; Table S1).

**Figure 1.**
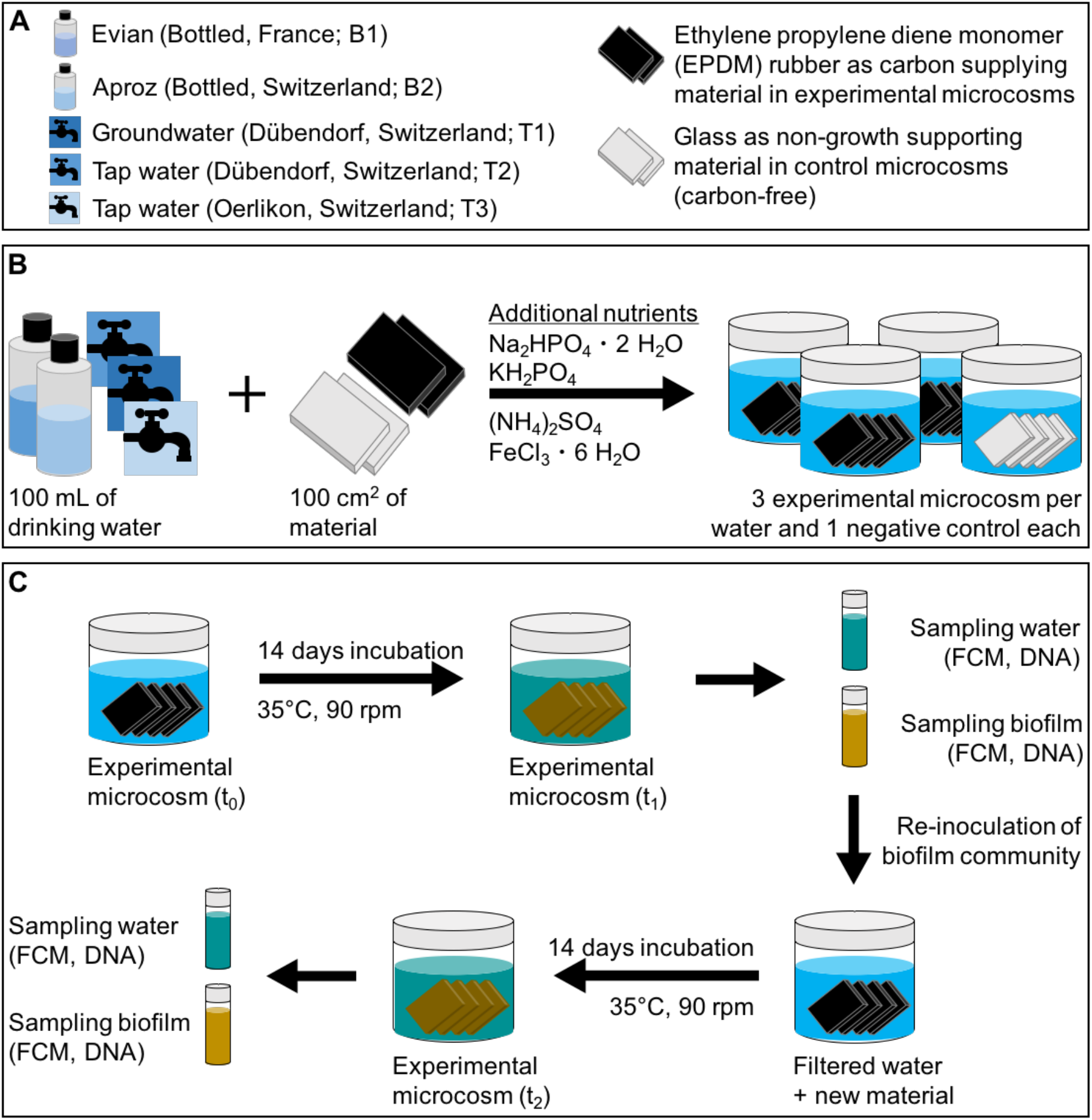
Experimental design. (A) Water and coupon materials. (B) Microcosms contained 100 mL water and 100 cm^2^ of coupons. To exclude growth limitation, phosphorous, nitrogen, and iron were added to each microcosm. (C) Microcosms were incubated for 14 days (35 °C, 90 rpm). Biofilms and water were analyzed using flow cytometry for total cell concentrations and 16S rRNA gene sequencing for the community compositions (t_1_). New microcosms were set up using filtered water which was spiked with individual biofilm communities. After another 14 days, biofilms and water phases were again sampled and analyzed (t_2_).

### 2.2 Microcosm design

Microcosms consisted of 240 mL glass jars (74 × 89 mm) with polypropylene lids including a PTFE lined inlet (Infochroma AG, Zug, Switzerland) (Figure 1). All glassware was cleaned with 1% (0.33 M) hydrochloric acid (HCl, 32%; Fluka, Sigma-Aldrich, Buchs, Switzerland), rinsed with nanopure water, and air dried. The clean glassware was muffled in a furnace (Nabertherm Schweiz AG, Hägendorf, Switzerland) (4.5 h; 500°C). EPDM flat sheets were cut into coupons of 0.2 × 2.6 × 4.3 cm (25 cm^2^). Prior to use, coupons were cleaned with a 0.1% (v/v) sodium hypochlorite solution (Sigma-Aldrich, Buchs, Switzerland) and rinsed with nanopure water. Glass was used as the control material. Microscope slides (Menzel-Gläser, 1 mm, ThermoScientific) were cut to the same coupon size as the EPDM and cleaned following the same procedure as for the glass jars (above). For each microcosm, four coupons or slides (i.e., 100 cm^2^) were stacked (Figure 1), separated by stainless steel springs. The springs and the jar lids were cleaned (60°C, 1 h) in a 100 g/L sodium persulfate solution (Na_2_S_2_O_8_, Sigma-Aldrich, Buchs, Switzerland), then rinsed with nanopure water, and air dried. Before use, the bottled drinking water was inverted 3-4 times for uniform mixing, while cold tap water was flushed for 5 min before filling into muffled 1 L SCHOTT Duran^®^ bottles (SCHOTT AG, Mainz, Germany). Each microcosm was subsequently filled with 100 mL water. To ensure unlimited growth conditions, additional nitrogen, phosphorous, and iron were added to the microcosms. The nitrogen/phosphorous buffer contained sodium phosphate dibasic dehydrate (Na_2_HPO_4_ • 2H_2_O, 1.28 g/L), potassium phosphate monobasic (KH_2_PO_4_, 0.3 g/L), and ammonium sulfate ((NH_4_)_2_SO_4_, 1.77 g/L) and 3.4 mL of buffer was added to each microcosm. Iron was supplemented in the form of iron (III) chloride hexahydrate (FeCl_3_ • 6 H_2_O, 2.7 g/L), with 50 μL per microcosm. All chemicals were purchased from Sigma-Aldrich (Buchs, Switzerland).

### 2.3 Migration and growth potential assays

For the assessment of carbon migration from the experimental material (EPDM) and the resulting consequences for bacterial growth, the material BioMig testing method was applied^18^. This method comprises a migration assay and a growth potential assay. In short: for the migration assay, 100 cm^2^ of EPDM was incubated (60°C, 24h, without shaking) with 100 mL bottled water (Evian). Over the course of seven days, the EPDM material was transferred into a new microcosm with fresh water every day. After the 1^st^, 3^rd^, and 7^th^ day of incubation, the water was sampled and the migrated total organic carbon (TOC) was quantified (see below). In addition, the growth of bacteria in the migration water was assessed. For this, 1 mL of fresh Evian bottled water was inoculated into 20 mL of migration water, with the addition of 690 μL phosphate/nitrogen buffer and 10 μL FeCl_3_. This test was performed in sterile, muffled 40 mL glass vials with screw caps lined with a PTFE septum (Supelco, Sigma-Aldrich Chemie GmbH, Buchs, Switzerland). Incubation (30°C, 120 rpm, 6 d) was followed by the quantification of the total cell concentration (TCC) using flow cytometry (FCM) (see 2.6.3). For the growth potential assay, 100 cm^2^ of new EPDM material was incubated (30°C, 14 d) with 100 mL of fresh bottled water (Evian) and additional nutrients (see 2.2). After 14 d of incubation, the water and biofilm phases were sampled for TCC, allowing for the determination of the bacterial growth potential within the experimental microcosms due to migrating carbon compounds (in direct comparison to a glass control set up without additional carbon)^18^.

### 2.4 Selection experiment

For all five water samples, triplicate microcosms were assembled with the testing material (EPDM) and an additional one containing glass as a control set up (Figure 1, B), as described above. After assembly (t_0_), the microcosms were incubated (14 d, 35°C, 90 rpm) (Figure 1, C). After 14 days (t_1_), biofilms were removed from the material surface (EPDM and glass; see 2.6.2) and both the biofilm and water phase of each microcosm were sampled for TCC (see 2.6.3) and community composition by 16S rRNA gene sequencing (see 2.6.4). For a second selection step, biofilm samples were re-inoculated into new microcosms. For this, the corresponding drinking water matrix was filtered using sterile bottle top filter units and membrane filters (Whatman^®^ Nucleopore™Track-Etched Membranes, 47 mm, 0.2 μm, Sigma Aldrich). New material was cleaned and stacked, additional nutrients added, and selected biofilm communities were added in a final concentration of 1 × 10^7^ cells/microcosm (i.e., 1 × 10^5^ cells/mL). After another 14d incubation, biofilms and water phases of all microcosms were again sampled (t_2_), following the same procedure. Regarding terminology, in the course of this study, initial drinking water communities are referred to as *original drinking water communities* and the biofilm communities of t_1_ and t_2_ as *selected biofilm communities*.

### 2.5 Attachment experiment

Here we compared attachment dynamics of selected biofilm communities with the original drinking water communities. The same microcosm set up was used as described above, with triplicate experimental microcosms (EPDM coupons) and single control microcosms (glass slides). The starting concentration of bacteria in the water phase (TCC, t_0_) was adjusted to be the same by diluting the biofilm communities relative to the drinking water TCC. The microcosms were incubated (35°C, 90 rpm) and the TCC in the water phase was measured for all at 30 min intervals over the course of 5 h (t_1_ – t_10_).

### 2.6 Sampling and analysis

#### 2.6.1 Chemical water analysis

Total organic carbon (TOC) was measured using a TOC-V_CPH_ Analyzer (Shimadzu Schweiz GmbH, Reinach, Switzerland). The minimum detection limit of the instrument is 0.1 mg/L. For total phosphorous, samples were chemically digested with potassium peroxodisulfate at 121°C, followed by a reaction to a phosphorous-molybdenum-blue complex and the determination of ortho-phosphate with spectrophotometry. The minimum detection limit of this method was 3.0 μg/L. Total nitrogen concentrations were measured via chemiluminescence using a Shimadzu TOC-L_CSH_ instrument. The minimum detection was 0.5 mg/L.

#### 2.6.2 Biofilm removal

All biofilms were removed with an electrical toothbrush (Oral-B^®^, Advanced Power) and toothbrush heads were replaced after each use to prevent cross-contamination. In short: EPDM or glass coupons were placed into muffled glass petri dishes and covered with filtered (0.2 μm) water. The water volume was always 25 mL per coupon (i.e., a total of 100 mL per microcosm). The coupons were brushed one by one, for approximately 90 sec each (including both coupon sides and the edges). During biofilm removal, 10 mL were saved and after the biofilm removal from all four coupons of a microcosm. This volume was ultimately used to recover biofilm residuals in the petri dish and on the brush head, by pouring the 10 mL filtered water into the petri dish and brushing without any coupon. The biofilm suspensions of the microcosms were collected in individual, sterile 100 mL SCHOTT Duran^®^ bottles. 10 mL of the biofilm suspension was used for flow cytometry (see 2.6.3). The rest of the biofilm suspension was used for the re-inoculation in the selection experiment (see 2.4), for further steps on community analysis (see 2.6.4), or for the attachment experiments (see 2.5).

#### 2.6.3 Flow cytometry for the quantification of total cell concentrations

FCM was used for the determination of total cell concentrations (TCC) in all biofilm and water samples. For biofilms, a 10 mL subsample of the biofilm suspension (see 2.6.3) was collected and needle-sonicated in a round-bottom glass tube (DURAN^®^; Faust Laborbedarf AG, Schaffhausen, Switzerland) using the Sonopuls HD 2200 instrument (Bandelin Sonorex, Rangendingen, Germany) and the Sonotrode Sonopuls MS 73 (tip diameter 3 mm, Bandelin). Sonication settings were: 30 sec at 50% power, and 40% amplitude intensity, with the pulse amplitude of the needle being 308 μm. The sonicated biofilm samples were then diluted 10-100x using 0.1 μm filtered Evian water (Millex^®^-VV, Merck-Millipore), while the water samples were measured undiluted. For the detection of TCC, samples were stained with 10 μL/mL SYBR^®^ Green I (SG, Invitrogen AG, Basel, Switzerland; 100x diluted in 10mM Tris buffer, pH 8). Stained samples were incubated (37°C, 10 min) and measured using a BD Accuri C6^®^ flow cytometer (BD, Belgium) or a CytoFLEX Flow Cytometer (Beckman Coulter International SA, Nyon, Switzerland). Gates and settings were kept the same within experiments. For more detailed information on data analysis and gating strategies see^19^.

#### 2.6.4 16S rRNA gene-based community analysis

For sequencing, samples of (1) all original drinking waters (t_0_), (2) all selected biofilms (t_1_, t_2_), and the water phase of the microcosms (t_1_, t_2_) were concentrated onto 0.2 μm polycarbonate Nucleopore^®^ membrane filters (ø 47 mm, Whatman, Kent, United Kingdom) by vacuum filtration, using sterile bottle top filter units. Filters were immediately frozen in liquid nitrogen and stored at −20°C.

#### 2.6.4.1 DNA extraction

The DNeasy PowerWater^®^ Kit (Qiagen, Hilden, Germany) was used for DNA extraction and performed according to the provided protocol. Extracted DNA was frozen and stored at −20°C until further processing.

#### 2.6.4.2 Library preparation and sequencing

For analyses on bacterial community compositions, the V3-V5 region of the 16S rRNA gene was amplified via polymerase chain reaction (PCR), using the primers Bakt_341F and Bakt_805R^20^. For library preparation, extracted DNA was quantified in duplicates using a Spark^®^ 10M Multimode Microplate Reader (Tecan, Switzerland; Qubit™ DNA Broad Range Assay). DNA concentrations were normalized between samples prior to amplification (1 ng DNA / 11 μL). For the PCR, normalized DNA was mixed with 2xKAPA HiFi HotStart Ready Mix (Kapa Biosystems, Roche Holding AG). Primers were added in a final concentration of 0.3 μM (details see Table S2, A). In addition to experimental samples, a negative control (i.e., amplification of sterile water instead of sample-DNA) and a positive control (self-made MOCK community: pure DNA of *Burkholderia xenovorans, Bacillus subtilus, Escherichia coli, Micrococcus luteus, Pseudomonas protegens, Paenibacillus sabinae*, and *Streptomyces violaceoruber*) were amplified. Additionally, some experimental samples were amplified in replicates. For this PCR, all samples were amplified in duplicates (2 × 25 μL reactions), which were combined prior to clean up. Amplified products were purified using the Agencort AMPure XO System (Beckman Coulter, Inc., Bera, CA, United States). For this, products were attached to magnetic beads, washed with 80% EtOH, and re-suspended in 10 mM Tris, pH 8.0. To enable pooling and re-identification of individual samples, specific sequencing Nextera XT v2 Index Kit adapters (Illumina) were annealed to the amplicons via Index PCR (Table S2, B). Products were again cleaned using the AMPure approach, quantified, and quality was checked using the High Sensitivity D1000 ScreenTape system (Agilent 2200 Tape Station). All samples were normalized to a concordant concentration followed by the pooling of 5 μL per sample. This pool was adapted to a final concentration of 2 mM and the base-pair (bp) length of the product determined with the Tape Station (627 bp).

Sequencing was performed using the MiSeq platform. For this, 0.1M NaOH was added to the pool, centrifuged (300 g, 60 s) and incubated for 5 min (room temperature) prior to the addition of the hybridization buffer HT1. This step was to (1) generate single stranded DNA and to (2) prevent unspecific bindings to the flow cell during sequencing. As a final step, 10% PhiX was added as a sequencing run control (Illumina: Technical Note on PhiX Control). The MiSeq run was a paired-end 600 cycle sequencing run. Data on community composition was generated in collaboration with the Genetic Diversity Center (GDC), ETH Zurich.

#### 2.6.4.3 Processing of sequencing data

Data processing followed a distinct pipeline. First, data quality was controlled using FastQC (Table S2, A). Then, read ends were trimmed and merged (Table S2, B), which was followed by an *in silico*-PCR and the trimming of the primer sites (Table S2, C). Finally, sequences were filtered (based on quality and size range) (Table S2, D) and amplicon sequence variants (ASV) were generated and taxonomically assigned. The clustering of sequences was performed as presented in a previous study^14^. It is based on an amplicon sequence variant (ASV) approach using UNOISE3, proposed by Edgar and colleagues^21^, and includes a correction for sequencing errors and a chimaeral removal. Clustered sequences are called zero-radius operational taxonomic units (ZOTUs). Due to a potential overestimation of the actual number of ZOTUs, an additionally clustering was performed at different identity levels of 99, 98, and 97%. For predictions on taxonomic assignments, the Silva 16S database (v128) and the SINTAX classifier were used (cut-off 0.9). See supplementary information for details on data analysis using R (Version 3.3.0) and RStudio (Version 1.1.477).

## 3. Results

### 3.1 Migration and growth potential assays

Applying an established material testing package (BioMig^18^) revealed that a considerable amount of organic carbon migrates from the experimental EPDM coupons and that a substantial fraction of the migrating carbon can be used by drinking water bacteria to grow. The migration assay (60 °C) showed that organic carbon migrated in high concentrations from new EPDM coupons and that it increased the TOC concentration of the water 5-fold (average: 1.1 μg TOC/mL, n = 2) within the first 24 h of migration (Figure 2A). The extent of migration diminished over time. However, even after 7 d of sequential migrations, the TOC concentration of the water with EPDM coupons still increased 3-fold (0.43 ± 0.03 μg TOC/mL, n = 2), equivalent to a rate of 0.3 μg TOC/cm^2^/d. These values are typical for flexible materials in contact with drinking water (e.g., in general^14,22^, or specifically for EPDM^23,24^). A separate growth potential assay at 30 °C showed that 2.3 ± 0.09 × 10^7^ cells/mL (n = 3) were able to grow on migrating carbon from EPDM coupons during 14 days, which is 30x more compared to growth in the absence of EPDM (Figure 2B). Given that the carbon-source for growth was the EPDM coupons, this translated to the growth of 2.3 × 10^7^ cells/cm^2^ coupon. Of these cells, 57 % (i.e., 1.3 × 10^7^ cells/cm^2^) were recovered directly from the surface of the EPDM coupons. To summarize, results show that the EPDM coupons favor biofilm formation by (1) providing a surface for colonization and by (2) adding biodegradable organic carbon to the water. Therefore, this material was deemed suitable for the further experiments on biofilm growth and the selection within growing communities (below).

**Figure 2.**
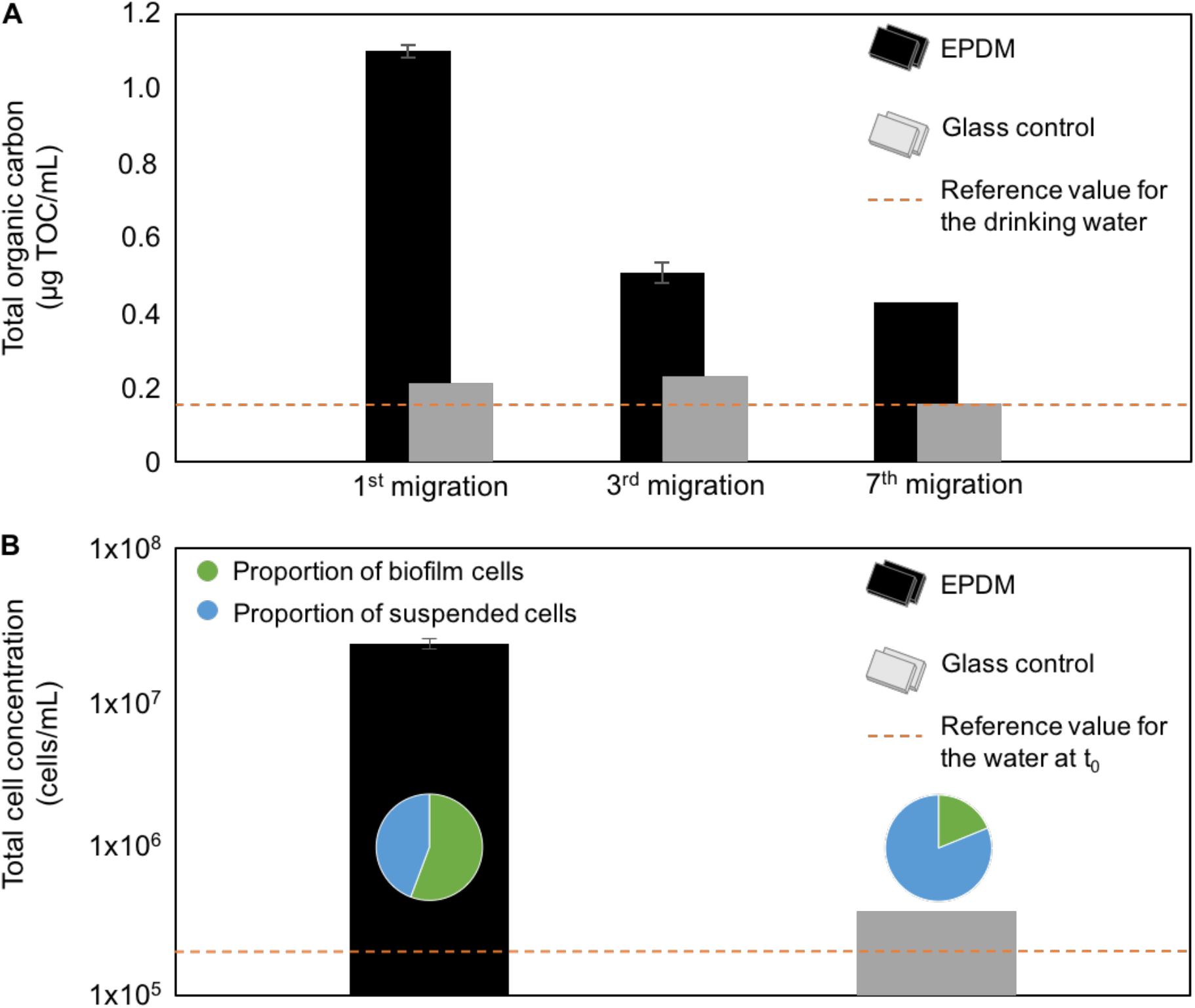
Assessment of carbon migration from EPDM and the resulting potential for bacterial growth. (A) Migration assay for the quantification of total organic carbon (TOC) in microcosms with EPDM or glass coupons. Material was transferred into fresh microcosms every 24 h, with measurements after the 1^st^, 3^rd^, and 7^th^ migration. (B) Bacterial growth potential based on EPDM, or glass as control. Total cell concentrations (TCC) are shown per mL for total growth of both suspended and biofilm cells. The conversion of cm^−2^ for biofilm cells to mL^−1^ was based on the water volume to material surface area ratio of 1:1 in the microcosm set up. Proportions of biofilm and suspended cells are indicated via pie charts. Error bars represent the range between duplicate microcosms in (A) and standard deviations for triplicate microcosms in (B).

### 3.2 Selection experiment

The basic concept and design of the growth potential assay was used to test the growth of five different drinking water communities on identical EPDM coupons (Figure 1). All communities showed (1) intensive growth and (2) a considerable loss in taxa diversity (apart from B2), with (3) the development of different biofilm communities involving shared taxa.

#### 3.2.1 Considerable growth for different original drinking water communities

Figure 3 shows that after two sequential 14-day cycles of inoculation and incubation (Figure 1), substantial growth was measured for all five waters in the presence of EPDM coupons, ranging within one order of magnitude (0.2 – 3.3 × 10^8^ cells/mL). These final concentrations represent both the planktonic and biofilm phases. The proportion of cells recovered directly from the biofilm ranged between 59 – 86 %, equivalent to 0.2 – 2 × 10^8^ cells/cm^2^. While experimental microcosms had considerable growth, differences were identified. Growth in the absence of EPDM coupons (i.e., in the glass controls) highlighted the impact of the migrating carbon, showing that TCC concentrations were 93 – 99 % lower without the additional carbon source. The proportion of cells in the biofilm was still high with 30 – 93 %, which translates to 1.4 ± 0.5 × 10^6^ cells/cm^2^ (n = 5). These findings confirm our results from Figure 2 on the carbon migration and growth potential based on EPDM coupons. The results show that the growth is high for different drinking water communities and that there was substantial biofilm growth, irrespective of the starting community.

**Figure 3.**
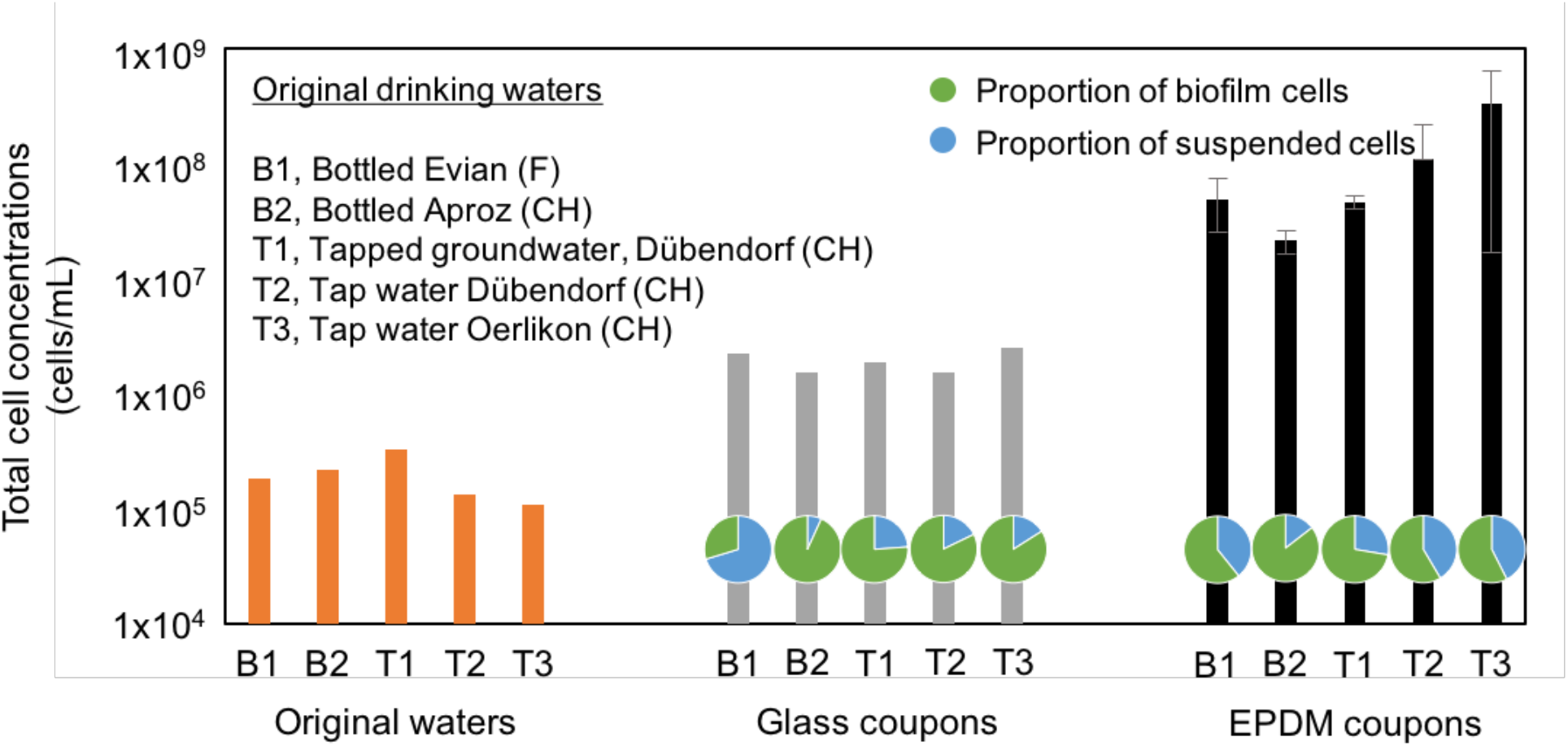
Total cell concentrations in the original drinking waters (t_0_) and in microcosms at the end of the experiment (t_2_), i.e., 2 × 14 d of incubation with an intermediate re-inoculation of biofilms grown at t_1_. For growth in microcosms with glass and EPDM coupons, total cell concentrations (TCC) are shown per mL for the total growth of both suspended and biofilm cells. The conversion of cm^−2^ for biofilm cells to mL^−1^ was based on the water volume to material surface area ratio of 1:1 in the microcosm set up. Proportions of biofilm and suspended cells are indicated via pie charts. Error bars represent standard deviations for triplicate microcosms in the EPDM coupon set up.

#### 3.2.2 Comparatively low taxa diversity detected in biofilm communities

A comparison of the original drinking water communities with the biofilm communities at the conclusion of the experiment revealed a notable loss in diversity (apart from B2, see below). Richness, i.e., the number of different taxa, decreased and became more comparable between the different waters. Figure 4A shows the richness values for the five original drinking water communities. Interestingly, tapped waters showed considerably more taxa (2’178 ± 131, n = 9) than the bottled waters, with the original water B2 comprising strikingly few taxa (54 ± 0, n = 3) compared to B1 (270 ± 15, n = 3). Overall, biofilm communities comprised comparatively few taxa (29 – 50 taxa), which corresponded to a diversity loss of 46 – 98 % from the original waters. This impressive loss of up to 2’000 individual taxa (tap waters) highlights the relevance of nutrient-based selection within establishing biofilm communities. As a consequence to this loss in diversity (through growth and selection), the similarity between the biofilm communities increased (with respect to diversity), with only 21 % variation in richness between biofilms as opposed to 73 % between the original drinking water communities. Shannon diversity followed a similar trend. Figure 4B shows a comparable dissimilarity between the original tapped water communities (5.9 ± 0.7, n = 9) and bottled water communities (2.3 ± 0.03, n = 3 for B1; 1.1 ± 0.01, n = 3 for B2). The relation between the Shannon Index (H’) and its maximum value (H’_max_) is important for drawing conclusions on diversity, i.e., the closer H’ to H’_max_, the higher the diversity within the community. Here, the relation was 1:1.3 for the tapped waters, 1:2.5 for B1, and 1:3.7 for B2 respectively, indicating a higher diversity in the tapped waters. This difference decreased with biofilm formation, resulting in a comparable degree of diversity. Here, the ratio between H’ and H’_max_ is close to 1:3 for all samples. This shows that (1) diversity decreased for (almost) all communities and (2) that biofilm communities are more similar to each other compared to the original drinking water communities. As indicated above, bottled water B2 was the misfit amongst the original water communities with a particularly low initial richness and diversity. Interestingly, this community also grew the least during the selection experiment (Figure 3). This suggests that the initially low diversity did not allow the community to metabolize as many nutrients as for the other more diverse communities. Results on richness and Shannon diversity allowed for the calculation of Evenness. Evenness indices were low for the bottled waters (0.34 ± 0.07, n = 6) and did not change much during the growth experiment (0.35 ± 0.09, n = 6). For the original tapped waters, Evenness was high (0.77 ± 0.08, n = 9), indicating a rather equal distribution of taxa. For the tapped water biofilms, indices decreased approx. 50 % which resulted in comparable Evenness indices for all samples (0.37 ± 0.1, n = 15).

**Figure 4.**
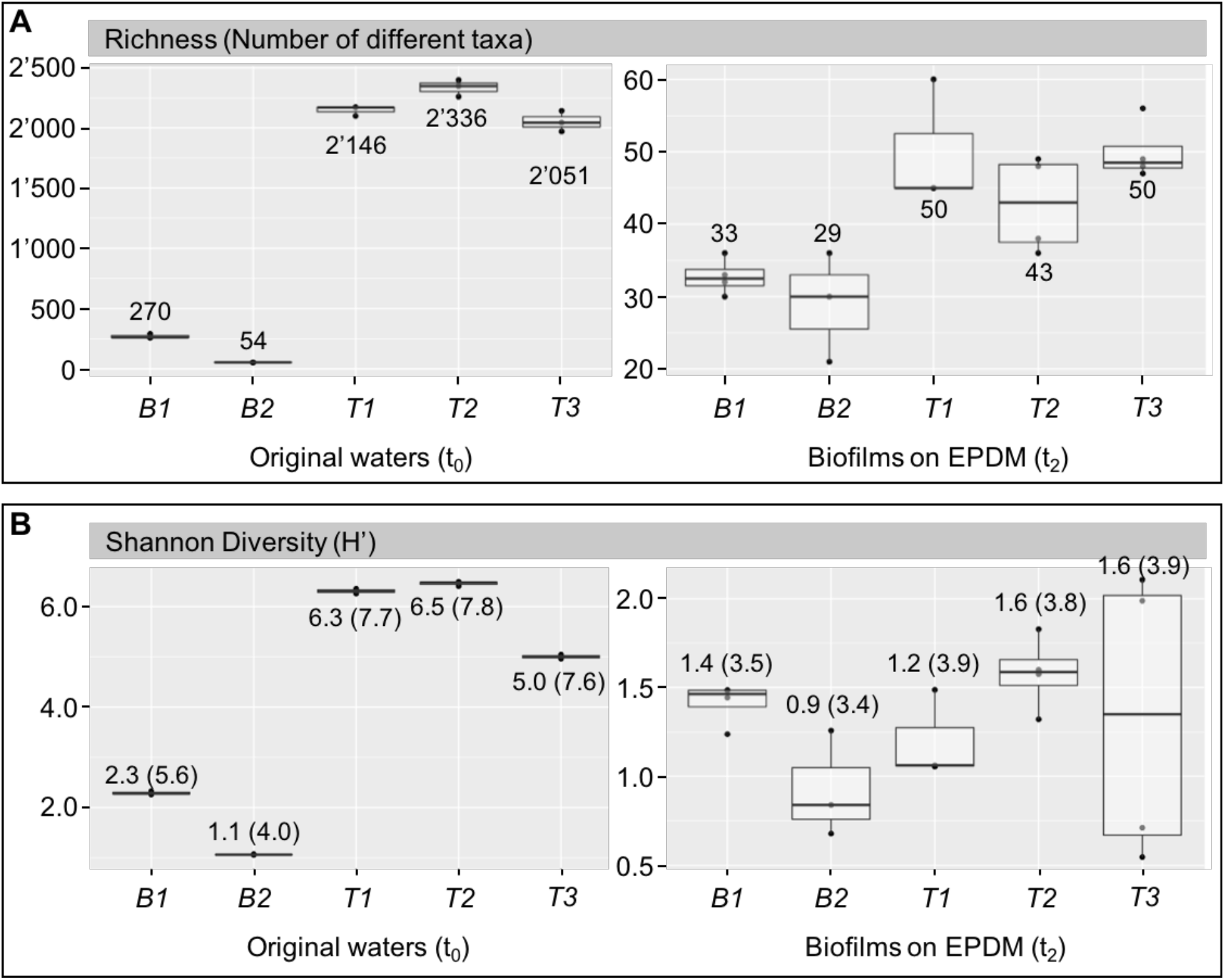
Diversity between microbial communities following growth on EPDM coupons. Alpha-diversity indices richness (A) and Shannon diversity (B) of the original drinking water communities (B1, B2, T1, T2, and T3) and the selected biofilm communities grown on EPDM (t_2_) for each experimental water. Results are presented as averaged numbers for richness (A) and averaged Shannon indices with additional information on (H’_max_). Original drinking waters were sequenced in triplicates and biofilms were sampled from triplicate experimental microcosms (additional sampling points in plot are due to replicate sampling of selected biofilms).

What is particularly interesting is that, involving already severe loss in taxa diversity and richness (i.e., a strong selection), the low degree of evenness within biofilm communities indicated a dominance of an even smaller number of taxa. This was, in fact, the case with the five most abundant taxa accounting for 95 ± 5 % (n = 15) of the individual biofilm community compositions (Table S4). These results highlight that (1) the bottled waters had low diversities from the start, (2) the number of different taxa decreased during biofilm formation and so did (3) the equality of their distribution. This rendered all biofilm communities more similar, with comparably low diversity and (4) only view taxa dominating the entire biofilm communities.

#### 3.2.3 Biofilm growth alters community composition

The decrease in taxa diversity came along with a change in community composition from the original drinking water to the selected biofilm communities. Figure 5 illustrates the dissimilarities between the bacterial communities of the original drinking waters (t_0_, triangles) and their corresponding biofilm communities that grew on EPDM coupons (t_2_, circles). The distance between original and biofilm communities was large for the tapped waters (e.g., highlighted for T3, Figure 5).

**Figure 5.**
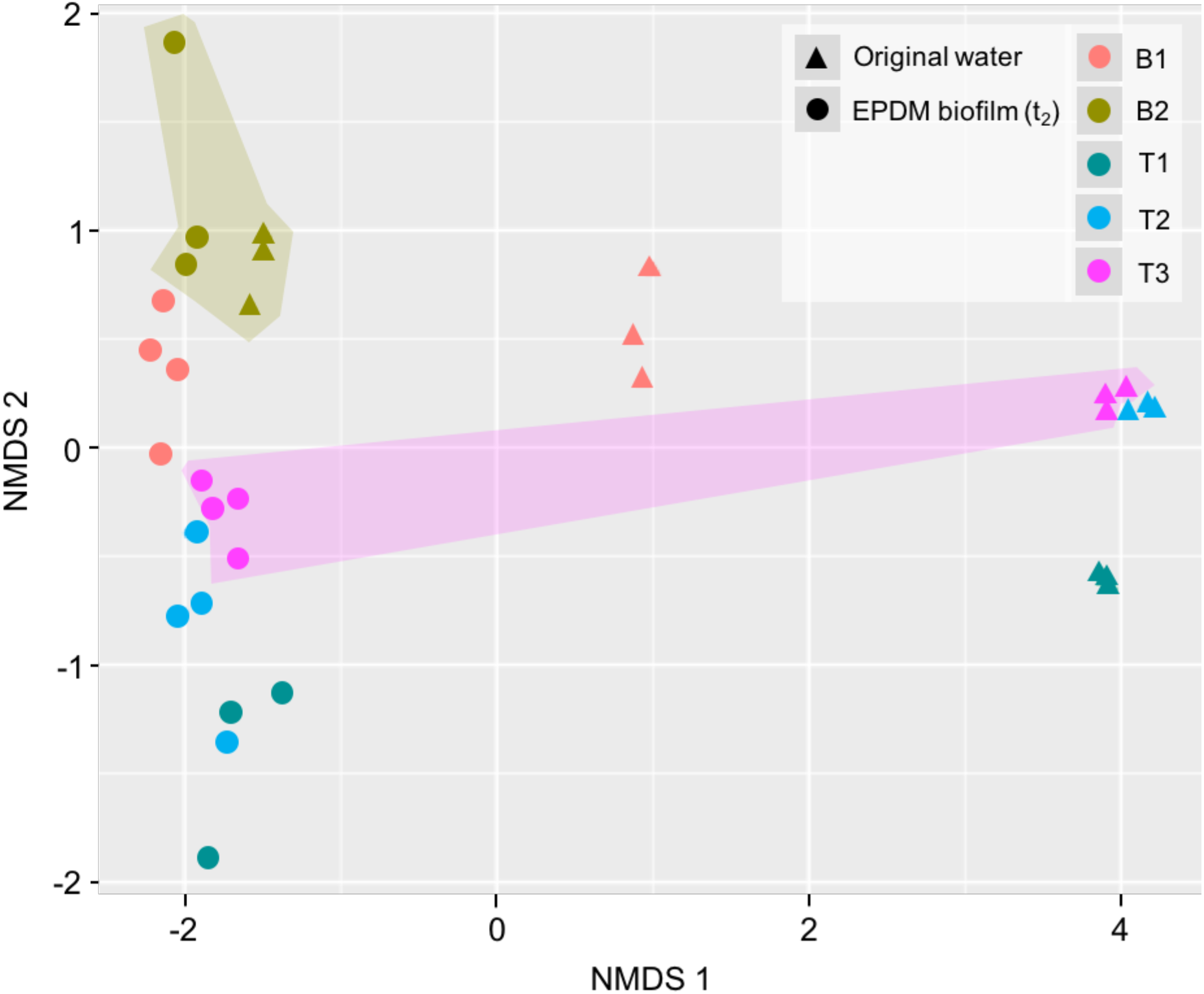
NMDS ordination plot based on Bray-Curtis Dissimilarity. Comparison of five different original drinking water communities (B1, B2, T1, T2, and T3) and their corresponding biofilms that formed on EPDM (t_2_). Original drinking waters were sequenced in triplicates and biofilms were sampled from triplicate experimental microcosms (additional sampling points in plot are due to replicate sequencing of selected biofilm samples). Highlighted areas were added manually to emphasize differences between samples.

In accordance with the degree of diversity loss, these community compositions changed considerably more compared to the bottled waters, which was attributed to the higher loss in taxa richness. Here, Bray-Curtis (BC) indices indicated a dissimilarity of 100 % between original and biofilm communities on taxa level (BC 1.0 ± 0.0, n = 3). In comparison, the dissimilarity was smaller for communities of B2 (highlighted in Figure 5), with BC indicating a dissimilarity of 89 %. The higher degree of similarity for B2 indicated that (1) the community composition changed comparatively little during biofilm formation. This indicates, in combination with the low growth, that diversity in the original B2 did not cover enough metabolic functions to exploit the full growth potential provided by migrating carbon. To emphasize this, 83 % of the most abundant taxa of B2 biofilms (i.e., the 5 most abundant taxa amongst triplicate microcosms) were also detected in the original water community (Table S5), which is high compared to B1 (56 %) or the tapped waters (38 – 50 %). Interestingly, when comparing samples of different origin (i.e., with different starting communities), biofilms were more similar to each other compared to the original drinking water communities. Here, Bray-Curtis dissimilarity between original drinking water communities was 0.94 ± 0.1 (n = 5) and resulted in a BC of 0.83 ± 0.16 (n = 5) between biofilms. In summary, accompanying the reduction in taxa diversity, the community compositions of the original waters changed during biofilm formation and become more similar to each other. What remains unclear is whether the loss in diversity and compositional changes necessarily involved the growth of identical taxa.

#### 3.2.4 Similarities in biofilm communities

Biofilm communities that developed from different (original drinking water) starting communities comprised shared taxa and families. Out of 29 ± 12 taxa/biofilm community (n = 15; including taxa with ≥ 0.01 % relative abundance), only two taxa were present in all biofilms that grew on EPDM coupons (Table S4). These taxa were identified as (1) *Methylobacterium* spp. (0.3 ± 0.1 %, n = 15; ZOTU7) and as (2) a not further identified member of the family Bradyrhizobiaceae (0.5 ± 0.6 %, n = 15; ZOTU23). Taxa that were detected in at least one of the experimental triplicates per set up were (1) a member of Bacilliaceae (0.04 ± 0.05 %, n = 13; ZOTU148), (2) *Aquabacterium* spp. (23 ± 29 %, n = 13; ZOTU1024), and (3) a member of the family Comamonadaceae (10 ± 16 %, n = 14; ZOTU5147). Here, ZOTU5147 was very abundant in the biofilms of B1 (35 ± 14 %, n = 3) and ZOTU1024 in the biofilms of T1 (70 ± 9 %, n = 3) and T2 (49 ± 8 %, n =3). Out of 16 ± 6 families/biofilm community (n = 15), four families were present in all EPDM biofilms, namely (1) Bradyrhizobiaceae (0.6 ± 0.6 %, n = 15), (2) Comamonadaceae (43 ± 28 %, n = 15), (3) Methylobacteriaceae (0.3 ± 0.1 %, n =15), and (4) Sphingomonadaceae (3 ± 7 %, n = 15) (Table S6). In addition, the families (1) Bacillaceae (0.05 ± 0.05 %, n = 13), (2) Brucellaceae (0.03 ± 0.02 %, n = 13), (3) Burkholderiaceae (22 ± 30 %, n = 13), (4) Caulobacteraceae (4 ± 6 %, n =13), and (5) Xanthomonadaceae (8 ± 13 %, n = 13) were present in at least one triplicate per experimental set up. Of these, Comamonadaceae was highly abundant in the biofilm communities of T1 (71 ± 10 %, n = 3) and T2 (61 ± 14 %, n = 3). For the bottled waters, Burkholderiaceae was dominant in B2 (71 ± 15 %, n = 3) and Xanthomonadaceae showed a high abundance in the biofilm communities of B1, with a relative abundance of up to 18 %. This result highlights that biofilm communities had a certain consistency in their compositions, despite clear differences in their starting communities. The selective pressure during biofilm formation and growth was not only demonstrated by the loss of taxa but also by the dominance of originally rare taxa. Within individual biofilm communities, the five most abundant taxa accounted for 94 ± 5 % (n = 15) (Table S4). From these individual abundant taxa, 53 ± 17 % (n = 5) were detectable in the corresponding original drinking water communities (i.e., the rest was below detection limit of the method). Interestingly, the chance of dominant biofilm taxa also being abundant in the original drinking water community was associated with the degree of initial diversity. For example, a highly abundant taxon in T1 biofilms (70 ± 10 %, n = 3; ZOTU1024) was rarely detected in the original water with a relative abundance of only 0.03 ± 0.01 % (n = 3) (Table S5). Bottled water B2 had the lowest taxa richness and diversity. Here, the most abundant biofilm taxa (71 ± 15 %, n = 3; ZOTU46) was already very abundant in the original water (8 ± 0.4 %, n = 3; ZOTU46). These results show that biofilm formation on carbon supplying EPDM coupons was highly selective and resulted in a considerable loss in taxa richness and diversity. As a result, the composition of biofilm communities differed from their original drinking water communities. Individual biofilms showed, however, similarities regarding dominant organisms, which indicated that, irrespective of starting communities, the environment (i.e., additional carbon supply) was selective for specific taxa and families, which was potentially linked to metabolic functions (see, e.g., ^25^).

### 3.3 Attachment experiment

The selected biofilm communities attached more and much faster to new surfaces compared to the original water communities (Figure 6).

**Figure 6.**
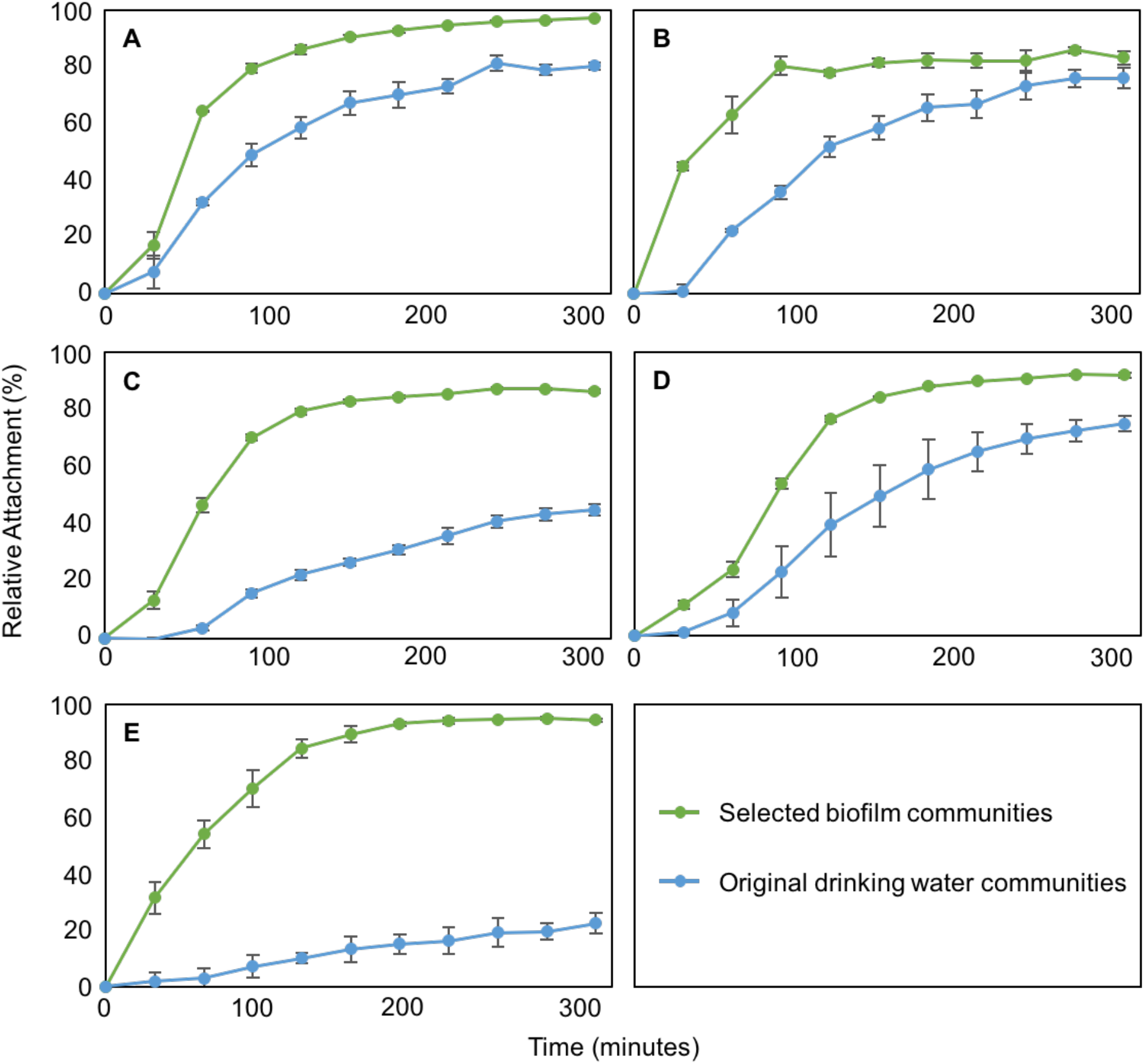
Attachment of selected biofilm and original drinking water communities to EPDM coupons. (A) B1 (bottled Evian; F), (B) B2 (bottled Aproz; CH), (C) T1 (tapped groundwater, Dübendorf; CH) (D) T2 (tap water, Dübendorf; CH) (E) T3 (tap water, Oerlikon; CH). Data points show average values and standard deviations for triplicate experiments.

For the selected communities, 91 ± 5 % (n = 15) of the cells attached within the first 5 hours after exposing freshly suspended cells to new EPDM coupons. This was approx. 30 % more than for the original water communities (69 ± 23 %, n = 15). Between the original communities, strong differences were measured in maximum attachment. For example, cells from original water B1 attached to 80 ± 1 % (n = 3) within 5 hours as opposed to T1 with only 45 ± 2 % (n = 3). The direct comparison between original and selected communities showed a clear advantage for the selected cells. For example, after 5 hours of incubation one tapped water (T3) showed a relative attachment of 94 ± 1 % (n = 3) for the selected community, but only 22 ± 4 % (n = 3) for the original community (Figure 6E). In absolute numbers, this percentage translates to a maximum attachment of 6.7 ± 0.1 × 10^4^ cells/cm^2^ (n = 3) for the selected community and 1.4 ± 0.3 × 10^4^ cells/cm^2^ (n = 3) for the original community (Table S6). In addition to the high maximum attachment, maximum attachment rates were on average 5.0 ± 1.7 × 10^4^ cells/cm^2^/h (n = 15) in selected communities and 2.4 ± 1.2 × 10^4^ cells/cm^2^/h (n = 15) in original communities. Regarding T3, maximum attachment rate for selected cells was almost 10-fold higher with 3.9 ± 0.4 × 10^4^ cells/cm^2^/h (n = 3) as opposed to 4.6 ± 0.9 × 10^3^ cells/cm^2^/h (n = 3). The comparison of the relative attachment between selected and original communities of all waters showed a 1- to 4-times higher maximum attachment and a 1- to 7-times higher maximum attachment rate for the selected communities. Interestingly, the attachment dynamics were similar with glass coupons as surface (Figure S1). Maximum attachment rates were almost identical between EPDM and glass coupons. Maximum attachment after 5 hours was, however, 6-fold higher on EPDM coupons (Table S7). In summary, a considerably large proportion of planktonic cells attached to the coupons (both EPDM and glass) within a short time. The selected communities attached faster and showed higher absolute values for attachment compared to the original drinking water communities.

## 4. Discussion

We analyzed biofilm growth on flexible EPDM coupons for five different drinking water communities (Figure 1). The purpose was to study the amount of growth due to biodegradable carbon migrating from the EPDM (Figure 2) and to assess selection within the developing biofilm communities due to this carbon. In the course of biofilm growth (Figure 3), all samples showed a significant loss in species diversity (Figure 4) with, however, the development of different community compositions (Figure 5). The selected communities were in turn more likely to form new biofilms on clean coupons (Figure 6).

### 4.1 Nutrient-based selection within microbial communities

Community composition in complex biofilms is governed by known ecological principles such as dispersal, selection, drift, and diversification^26^. In the presence of sufficient nutrients, selection within microbial communities is (at least partially) driven by the metabolic potential and growth physiology of individual members. The supply of different/new substrates to an established community allows the growth of different bacterial species based on their metabolic capabilities, resulting in a change in the community composition. For example, Eilers and colleagues^27^ (soil communities) and Reintjes and colleagues^28^ (marine communities) demonstrated that the addition of particular carbon substrates resulted in bacterial growth, a loss in diversity, and considerable changes in community compositions. Reintjes and colleagues^28^ also showed that initially abundant taxa were not abundant in the grown communities anymore. Finally, Wawrik and colleages^29^ showed that carbon sources that differ in their complexity select for different bacterial communities, with 70 % dissimilarities between communities grown on different substrates. This happened quickly, e.g., with a developing community growing on acetate being 70 % dissimilar within 18 h of incubation. The establishment of similar communities, relative to the complexity of supplied carbon substrate, indicated that metabolic capabilities (biochemical pathways) are similar amongst growing cells^29^. Our results mirrored many of these findings, in that richness decreased dramatically (Figure 4) and the dominating bacteria after growth on EPDM were completely different to the dominating bacteria in each original water (Figure 5). However, it is noted that even when microorganisms are capable of using the same substrates, growth physiology (i.e., growth rate and yield) allows some species to outcompete others. This so called *competitive exclusion principle*^30^ was demonstrated by Friedman and colleagues^31^ and Christensen and colleagues^32^ who correlated the ability to outcompete others to a species’ growth rate and yield. Our data (e.g., Figure 5) does not allow separation between selection caused by metabolic capabilities and growth physiology. However, this may be an explanation why some taxa dominated in the microcosms.

The arguments above explain selection during growth, but may lead to an erroneous conclusion that growth on the EPDM coupons should by default result in similar communities being selected. Here, our data clearly showed that all five original water samples resulted in completely different final communities following growth (Figure 5). This is explained by the fact that many different bacterial species can have the same carbon-degrading functions, i.e., identity does not equal functionality. In this regard, Burke and colleagues^33^ showed that communities that are dissimilar in their taxonomical composition (e.g., 15 % similarity) can be very similar regarding their functional composition (with, e.g., 70 % similarity). Recent work by Goldford and colleagues^25^ showed that growth on additional carbon sources increased the dissimilarity between developing communities on taxonomic level, however, revealed carbon source specific analogies on family level^25^.

### 4.2 Understanding building plumbing microbiomes

One very practical relevance of biofilm growth and selection discussed above is the first colonization of drinking water plumbing systems during the commissioning of new buildings. Here, a wide variety of synthetic plumbing materials^34,35^ provides biodegradable organic carbon^36,37^ to complex drinking water microbial communities^38,39^. The consequence is biofilm formation and development in the plumbing system, which ultimately affects the microbiological quality of the drinking water^34,40^. Observations in recent years from several pilot-scale and full-scale studies reported considerable changes in drinking water microbial communities after passage through building plumbing systems. For example, Ling and colleagues^11^ showed that the community composition of the drinking water changes during stagnation within building plumbing systems compared to the composition within the distribution network. Work from our own group specifically compared tap water, stagnated water, and biofilm communities in shower hoses and showed distinct changes in the microbiome due to the biofilms growing in the hoses^12^. These findings are further supported by data from Ji and colleagues^13^ and Dai and colleagues^41^ showing that material, temperature, and stagnation time change the microbiology compared to the water community flowing into the rig installations, with, e.g., stagnation resulting in a diversity loss within the drinking water community^41^.

It is clear that building plumbing systems are per se complex environments, with multiple confounding factors (e.g., temperature, hydraulics, nutrients) affecting bacterial colonization, growth, and microbiome composition. Previous studies suggested that the choice of plumbing material plays a critical role, particularly when the material supplies nutrients for growth^15,24,42^. For example, Rogers and colleagues^43^ studied biofilm development on different materials with different extents of growth supporting properties (and their ability to resist invading *Legionella*). Proctor and colleagues^15^ studied different shower hose materials and found differences in growth community composition. Along the same lines, we demonstrated in a previous study considerable selection in biofilm communities forming within shower hoses, with differences in the microbiome measurable on small-scale^14^. In the present study, we show an example for EPDM, which is commonly used for drinking water applications^16,24,44,45^ (Figures 3, 4, 5). We demonstrated selection, but also showed that selection differed based on the source water (Figure 5). While there is an obvious need and scope for larger observational studies on drinking water microbiomes^46^, there is also a clear need for basic laboratory-scale ecological studies that can help to inform on interpretations from complex building plumbing data. Moreover, understanding the basic ecology of building plumbing systems will provide a basis for proactive management of the microbiomes in these systems.

### 4.3 Managing colonization of building plumbing materials

Better knowledge on growth-dependent selection within biofilm communities can be used to design building plumbing systems where the microbiology is controlled or even specifically tailored to the system. Microbial colonization and growth on building plumbing materials is currently not (properly) controlled. Upon commissioning of a building, all new plumbing material is exposed to complex drinking water communities during the first use. In fact, there is essentially no control over the identity and composition of bacteria that attach and proliferate in the new system, irrespective of the location, source water, disinfectant use, building type, or plumbing materials. To date, there are surprisingly few studies looking at this initial colonization of building plumbing materials, both full- and pilot-scale. A notable exception is the study by Salehi and colleagues^34^, where they monitored changes in water chemistry and bacterial growth during the first days/weeks of building occupation. Also, a study by Douterelo and colleagues^47^, showed that specific bacteria are dominant during the initial colonization (7 – 28 d) of distribution pipe materials. The fact is, in current practice the owners/operators have effectively no control over the communities that colonize their building plumbing systems.

Smart use of material properties can control microbial growth (and thus biofilm communities). For example, the use of high quality materials and the avoidance of low quality ones (e.g., flexible hoses) reduces the potential of bacteria to actually grow. For this, standards for material quality requirements have already been implemented in Europe (e.g., ^48^) and official tests on carbon migration and corresponding growth potentials have been established (e.g., ^17,49^) (e.g., Figure 2). Using such tests to qualify the use of individual materials in new buildings should be a must for the industry. A more expensive but sensible approach is to use materials that do not leach any carbon (e.g., stainless steel plumbing). For example, Van der Kooij and colleagues^50^ showed that biofilm growth and the incorporation of *Legionella* spp. was less on stainless steel compared to polymeric PE-X pipes. It is, however, important to take into account that high quality polymeric materials can perform as good as metal piping with regard to microbial growth (see, e.g., ^51^). As another possibility, some studies suggested developing and using plumbing materials with anti-microbial properties for minimizing microbial growth and the proliferation of pathogens. For example, Saleh and colleagues^52^ showed that a coating containing copper and silver ions resulted in less bacterial attachment and biofilm growth when exposing a *Pseudomonas aeruginosa* isolate (10^5^ – 10^6^ CFU/mL) to coated glass slides for 2 h, shaking.

An alternative approach could be to embrace the microbiology of building plumbing systems instead of resisting it. Carbon migration from building plumbing materials can theoretically be used to select and maintain preferred communities. Wang and colleagues^53^ provocatively suggested that systems may be redesigned in a pre/probiotic approach to favor certain communities of choice. One way to approach this addresses the concept of niche occupation, which is especially important during the colonization of new surfaces, e.g., during the commissioning of a new building. Niche occupation can result in the exclusion of species due to a more efficient spatial expansion of a competitor or due to better growth physiologies^54^. For the first, Schluter and colleagues^55^ emphasized the importance of adhesion during initial attachment for the *evolutionary fate of microbes in biofilms*. For the second, Freilich and colleagues^56^ defined the competition for identical nutrient sources as a *win-lose relationship*, which will ultimately allow organisms with better growth yields/rates to outcompete others. This pre/probiotic approach can be taken a step further by introducing, selecting, and maintaining specific antagonistic bacteria that challenge unwanted organisms. For example, several studies showed that a range of aquatic isolates, especially *Pseudomonas* spp., produce *bacteriocin-like substances* that have an antagonistic effect on the establishment of *Legionella* spp. in biofilms^57–59^, which can potentially be exploited as probiotic communities against *Legionella pneumophila*.

Here we propose a combination of the approaches above. We argue for the use of plumbing materials that provide specific substrates and for the targeted colonization of these materials of a benign microbial community. The approach foresees the use of materials that migrate organic carbon in such a quality and quantity that it allows bacteria to grow and to sustain their existence in the developing biofilm. We furthermore propose colonizing these materials with bacteria from a safe source (e.g., bottled water), pre-selected on the substances migrating from the material (e.g., Figure 5). This adaption to the nutrients ultimately allows for a rapid colonization (e.g., Figure 6), growth and long term persistence. A further expansion of the approach could be the use of purposefully designed synthetic communities that specifically include antagonists to specific building plumbing pathogens^57–59^. Combining both niche occupancy capabilities and powerful antagonistic functions within a pre-conditioned/pre-selected community is an unconventional but exciting approach towards the future management of biofilm formation on polymeric materials in contact with drinking water.

## 5. Conclusions

- The use of a flexible polymeric plumbing material (here EPDM) increased the biodegradable organic carbon concentration of drinking water, which resulted in substantial growth for bacterial communities of different origin.
- The migrating carbon drove nutrient-based selection within the original drinking water communities, which resulted in (1) a dramatic decrease in taxa richness and diversity, (2) compositional changes in communities, and (3) an increase in similarity amongst growing biofilm communities, i.e., similarities in abundant taxa.
- Selected biofilm bacteria showed better attachment performances to new material surfaces, with more attachment and higher attachment rates.
- This work is a step towards pro-active managing of building plumbing biofilms through nutrient-based selection of specific communities of choice.

## Acknowledgements

The authors thank Silvia Kobel and Aria Minder-Pfyl for support and protocols for library preparation and sequencing, Jean-Claude Walser for support with data processing, and Caitlin Proctor for preliminary discussions on the topic.

## Funding

This work was funded by the Swiss National Science Foundation (SNSF grant nr. 31003A_163366/1).

## Author contributions

LN: experimental design, experimental work, data analysis, and manuscript writing.

LC: experimental design, experimental work, and data analysis.

FH: experimental design and manuscript writing.

## Notes

### Competing Interest Statement

The authors have declared no competing interest.

