## Supplementary information for "Towards a probiotic approach for building plumbing – nutrient-based selection during initial biofilm formation on flexible polymeric materials"

### Table of contents

**Table S1** Chemical and biological composition of experimental drinking waters

**Table S2** Settings for Amplification PCR and Index PCR reactions

**Table S3** Report information on data processing

**Table S4** Relative abundances of taxa within biofilm communities grown on EPDM coupons.

**Table S5** Shared taxa between biofilm communities and their original waters

**Table S6** Relative abundances of families within biofilm communities grown on EPDM coupons.

**Table S7** Attachment performances of selected original communities onto EPDM or glass coupons.

**Figure S1** Attachment of selected and original communities onto glass coupons.

**Table S1** Chemical and biological composition of different drinking waters used in this study. Total phosphorous, total nitrogen, total organic carbon, and total bacterial numbers are displayed for (1) Bottled Evian (France, B1), (2) Bottled Aproz (Switzerland, B2), (3) tapped Groundwater Dübendorf (Switzerland, T1), (4) Tap water Dübendorf (Switzerland, T2), and (5) Tap water Oerlikon (Switzerland, T3).

| | **Total phosphorous (µg/mL)** | **Total nitrogen (mg/mL)** | **Total organic carbon (mg/mL)** | **Bacterial numbers (TCC/mL)** |
|---|---|---|---|---|
| **Evian (Bottled, F)** | 2.4 | 0.9 | 0.08 | $1.8 \times 10^5$ |
| **Aproz (Bottled (CH)** | 5.3 | <0.5 | 0.19 | $2.2 \times 10^5$ |
| **Groundwater (Dübendorf, CH)** | 23.5 | 3.0 | 1.09 | $3.3 \times 10^5$ |
| **Tap water (Dübendorf, CH)** | 7.8 | 1.8 | 0.44 | $1.4 \times 10^5$ |
| **Tap water (Oerlikon, CH)** | 6.9 | 0.7 | 0.37 | $1.1 \times 10^5$ |

**Table S2** Information on volumes for amplifications and on settings of (A) amplification PCR and (B) Index PCR reactions.

| | | |
|---|---|---|
| **(A) Amplicon PCR** | | |
| | | *Volume (25 µL reaction)* |
| 2xKAPA HiFi HotStart ReadyMix | | 12.5 µL |
| Forward primer (10 µM) | | 0.75 µL |
| Reverse primer (10 µM) | | 0.75 µL |
| Template DNA (adjusted to 1 ng / reaction with DNase free water) | | 11.0 µL |
| *Temperature* | *Duration* | *Cycles* |
| 95 °C | 5:00 min | |
| 95 °C | 0:20 min | 29 x |
| 51 °C | 0:15 min | |
| 72 °C | 0:30 min | |
| 4°C | hold | |
| **(B) Index PCR** | | |
| | | *Volume (50 µL reaction)* |
| 2xKAPA HotStart ReadyMix | | 25.0 µL |
| Nextera XT Index 1 primer | | 5.0 µL |
| Nextera XT Index 2 primer | | 5.0 µL |
| Template DNA | | 15.0 µL |
| *Temperature* | *Duration* | *Cycles* |
| 95 °C | 3:00 min | |
| 95 °C | 0:30 min | 8 x |
| 51 °C | 0:35 min | |
| 72 °C | 0:30 min | |
| 4°C | hold | |

**Table S3** Report information of data processing.

| | | |
|---|---|---|
| **(A) Quality control** | | |
| | FastQC | |
| **(B) Trimming and merging of primers** | | |
| | usearch | v11.0.667_i86linux64 |
| | Trim R1 | 20 |
| | Trim R2 | 40 |
| | **Flash** | v1.2.11 |
| | Minimal overlap | 15 |
| | Maximal overlap | 300 |
| | Maximal mismatch density | 0.25 |
| **(C) Primer site trimming** | | |
| | **Usearch** | v11.0.667_ i86linux64 |
| | Coverage | full-length |
| | Allowed number of mismatches | 3 |
| | Amplicon size range | 100 - 600 |
| **(D) Filtering based on quality and size** | | |
| | Size range | 200 - 500 |
| | GC range | 30 - 70 |
| | Minimal Q mean | 20 |
| | Number of Ns | 0 |
| | Low complexity | dust / 30 |

**Table S4** Relative abundances of ZOTUs (taxa) within biofilm communities that formed on EPDM coupons and with different original starting communities (B1, B2, T1, T2, T3).

| | Biofilm 1 | | Biofilm 2 | | Biofilm 3 | |
|---|---|---|---|---|---|---|
| | **Rel %** | **ZOTU** | **Rel %** | **ZOTU** | **Rel %** | **ZOTU** |
| **B1** | 49.6 | ZOTU10 | 42.2 | ZOTU1024 | 51.6 | ZOTU5147 |
| | 29.3 | ZOTU46 | 36.8 | ZOTU5147 | 14.2 | ZOTU35 |
| | 16.6 | ZOTU5147 | 6.4 | ZOTU35 | 12.4 | ZOTU10 |
| | 1.6 | ZOTU61 | 4.2 | ZOTU2255 | 10.8 | ZOTU46 |
| | 0.8 | ZOTU9849 | 3.5 | ZOTU21 | 8.1 | ZOTU2255 |
| | 0.4 | ZOTU5674 | 2.2 | ZOTU10 | 0.7 | ZOTU61 |
| | 0.3 | ZOTU1816 | 1.4 | ZOTU115 | 0.7 | ZOTU141 |
| | 0.3 | ZOTU141 | 1.2 | ZOTU444 | 0.6 | ZOTU77 |
| | 0.3 | ZOTU77 | 0.6 | ZOTU11 | 0.3 | ZOTU7 |
| | 0.2 | ZOTU729 | 0.5 | ZOTU141 | 0.2 | ZOTU115 |
| | 0.2 | ZOTU7 | 0.3 | ZOTU46 | 0.1 | ZOTU1816 |
| | 0.1 | ZOTU23 | 0.2 | ZOTU11189 | 0.1 | ZOTU23 |
| | 0.08 | ZOTU421 | 0.2 | ZOTU7 | 0.06 | ZOTU11380 |
| | 0.04 | ZOTU148 | 0.07 | ZOTU23 | 0.04 | ZOTU142 |
| | 0.03 | ZOTU42 | 0.07 | ZOTU61 | 0.03 | ZOTU148 |
| | 0.03 | ZOTU22 | 0.04 | ZOTU142 | 0.03 | ZOTU5674 |
| | 0.03 | ZOTU2436 | 0.04 | ZOTU77 | 0.02 | ZOTU13679 |
| | 0.02 | ZOTU35 | 0.03 | ZOTU4844 | 0.02 | ZOTU4 |
| | 0.02 | ZOTU3965 | 0.02 | ZOTU1816 | 0.01 | ZOTU42 |
| | 0.01 | ZOTU13679 | 0.02 | ZOTU2436 | 0.01 | ZOTU113 |
| | 0.01 | ZOTU113 | 0.02 | ZOTU5674 | 0.01 | ZOTU2436 |
| | 0.01 | ZOTU6380 | 0.02 | ZOTU4 | 0.01 | ZOTU6380 |
| | 0.01 | ZOTU7519 | 0.02 | ZOTU148 | 0.01 | ZOTU5438 |
| | 0.01 | ZOTU21 | 0.01 | ZOTU6380 | 0.01 | ZOTU3965 |
| | | | 0.01 | ZOTU42 | 0.01 | ZOTU1024 |
| | | | 0.01 | ZOTU113 | 0.01 | ZOTU21 |
| | | | 0.01 | ZOTU4269 | | |
| | | | 0.01 | ZOTU11380 | | |
| | | | 0.01 | ZOTU144 | | |
| | | | 0.01 | ZOTU778 | | |
| **B2** | 84.6 | ZOTU46 | 78.5 | ZOTU46 | 49.3 | ZOTU46 |
| | 5.5 | ZOTU21 | 9.0 | ZOTU5438 | 32.8 | ZOTU5438 |
| | 4.3 | ZOTU31 | 5.7 | ZOTU21 | 10.2 | ZOTU21 |
| | 2.6 | ZOTU5438 | 3.2 | ZOTU31 | 4.0 | ZOTU5138 |
| | 1.3 | ZOTU5138 | 1.8 | ZOTU77 | 1.5 | ZOTU77 |
| | 0.4 | ZOTU7 | 1.3 | ZOTU5138 | 1.0 | ZOTU421 |
| | 0.3 | ZOTU77 | 0.1 | ZOTU421 | 0.6 | ZOTU31 |
| | 0.3 | ZOTU5674 | 0.1 | ZOTU7 | 0.2 | ZOTU7 |
| | 0.2 | ZOTU87 | 0.05 | ZOTU2052 | 0.2 | ZOTU2052 |
| | 0.1 | ZOTU23 | 0.05 | ZOTU23 | 0.09 | ZOTU23 |
| | 0.07 | ZOTU2207 | 0.02 | ZOTU5725 | 0.07 | ZOTU5147 |
| | 0.03 | ZOTU148 | 0.02 | ZOTU5674 | 0.02 | ZOTU42 |
| | 0.02 | ZOTU421 | 0.02 | ZOTU113 | 0.02 | ZOTU2436 |
| | 0.02 | ZOTU42 | 0.01 | ZOTU4 | 0.02 | ZOTU148 |
| | 0.02 | ZOTU113 | 0.01 | ZOTU1024 | 0.01 | ZOTU6117 |
| | 0.02 | ZOTU2436 | 0.01 | ZOTU87 | 0.01 | ZOTU3965 |
| | 0.02 | ZOTU4311 | | | 0.01 | ZOTU5 |

| | | | | | | |
|---|---|---|---|---|---|---|
| | 0.02 | ZOTU4498 | | | 0.01 | ZOTU20 |
| | 0.01 | ZOTU4295 | | | 0.01 | ZOTU113 |
| | 0.01 | ZOTU6117 | | | 0.01 | ZOTU28 |
| | **Biofilm 1** | | **Biofilm 2** | | **Biofilm 3** | |
| | **Rel %** | **ZOTU** | **Rel %** | **ZOTU** | **Rel %** | **ZOTU** |
| | 0.01 | ZOTU2052 | | | 0.01 | ZOTU7365 |
| | 0.01 | ZOTU9847 | | | 0.01 | ZOTU10 |
| | 0.01 | ZOTU35 | | | 0.01 | ZOTU6049 |
| | 0.01 | ZOTU4599 | | | 0.01 | ZOTU4599 |
| | 0.01 | ZOTU4 | | | | |
| | 0.01 | ZOTU11702 | | | | |
| | 0.01 | ZOTU7365 | | | | |
| | 0.01 | ZOTU2605 | | | | |
| | 0.01 | ZOTU165 | | | | |
| **T1** | 56.7 | ZOTU1024 | 76.9 | ZOTU1024 | 76.8 | ZOTU1024 |
| | 15.2 | ZOTU34 | 5.0 | ZOTU25 | 5.3 | ZOTU13 |
| | 13.5 | ZOTU9139 | 4.0 | ZOTU94 | 4.3 | ZOTU25 |
| | 3.9 | ZOTU13 | 3.5 | ZOTU185 | 3.2 | ZOTU4295 |
| | 3.1 | ZOTU4295 | 3.4 | ZOTU4295 | 2.6 | ZOTU94 |
| | 1.5 | ZOTU2448 | 2.0 | ZOTU13 | 1.8 | ZOTU64 |
| | 1.4 | ZOTU64 | 1.3 | ZOTU5147 | 1.3 | ZOTU185 |
| | 1.1 | ZOTU23 | 1.0 | ZOTU23 | 1.1 | ZOTU32 |
| | 0.7 | ZOTU25 | 0.5 | ZOTU12603 | 0.9 | ZOTU2448 |
| | 0.5 | ZOTU32 | 0.5 | ZOTU7874 | 0.9 | ZOTU23 |
| | 0.5 | ZOTU185 | 0.4 | ZOTU5 | 0.8 | ZOTU157 |
| | 0.4 | ZOTU7 | 0.2 | ZOTU2448 | 0.2 | ZOTU7874 |
| | 0.2 | ZOTU12603 | 0.2 | ZOTU32 | 0.2 | ZOTU7 |
| | 0.2 | ZOTU157 | 0.2 | ZOTU7 | 0.1 | ZOTU12603 |
| | 0.2 | ZOTU7874 | 0.1 | ZOTU151 | 0.06 | ZOTU160 |
| | 0.2 | ZOTU151 | 0.1 | ZOTU444 | 0.04 | ZOTU151 |
| | 0.1 | ZOTU2881 | 0.1 | ZOTU5329 | 0.04 | ZOTU13720 |
| | 0.1 | ZOTU160 | 0.08 | ZOTU10854 | 0.03 | ZOTU2436 |
| | 0.09 | ZOTU5147 | 0.07 | ZOTU160 | 0.03 | ZOTU5329 |
| | 0.08 | ZOTU94 | 0.05 | ZOTU13720 | 0.02 | ZOTU148 |
| | 0.07 | ZOTU5329 | 0.03 | ZOTU578 | 0.02 | ZOTU43 |
| | 0.07 | ZOTU148 | 0.03 | ZOTU2881 | 0.02 | ZOTU5147 |
| | 0.04 | ZOTU220 | 0.03 | ZOTU141 | 0.02 | ZOTU2881 |
| | 0.03 | ZOTU42 | 0.02 | ZOTU220 | 0.01 | ZOTU220 |
| | 0.02 | ZOTU578 | 0.02 | ZOTU113 | 0.01 | ZOTU5725 |
| | 0.02 | ZOTU7300 | 0.01 | ZOTU7300 | 0.01 | ZOTU42 |
| | 0.02 | ZOTU13720 | 0.01 | ZOTU145 | 0.01 | ZOTU10854 |
| | 0.02 | ZOTU2436 | 0.01 | ZOTU7347 | 0.01 | ZOTU113 |
| | 0.01 | ZOTU3179 | 0.01 | ZOTU43 | 0.01 | ZOTU13584 |
| | 0.01 | ZOTU444 | 0.01 | ZOTU64 | 0.01 | ZOTU5 |
| | 0.01 | ZOTU113 | 0.01 | ZOTU6119 | 0.01 | ZOTU4918 |
| | 0.01 | ZOTU5892 | 0.01 | ZOTU11348 | 0.01 | ZOTU11348 |
| | 0.01 | ZOTU10854 | 0.01 | ZOTU10190 | 0.01 | ZOTU5892 |
| | 0.01 | ZOTU4498 | 0.01 | ZOTU5892 | 0.01 | ZOTU578 |
| | 0.01 | ZOTU43 | 0.01 | ZOTU307 | 0.01 | ZOTU7745 |
| | 0.01 | ZOTU3344 | 0.01 | ZOTU5725 | 0.01 | ZOTU9139 |
| | 0.01 | ZOTU4546 | 0.01 | ZOTU933 | 0.01 | ZOTU125 |
| | 0.01 | ZOTU144 | 0.01 | ZOTU31 | | |
| | 0.01 | ZOTU3802 | 0.01 | ZOTU2628 | | |
| | 0.01 | ZOTU82 | | | | |
| | 0.01 | ZOTU3949 | | | | |
| | 0.01 | ZOTU146 | | | | |

| | | | | | | |
|---|---|---|---|---|---|---|
| | 0.01 | ZOTU202 | | | | |
| | 0.01 | ZOTU46 | | | | |
| | 0.01 | ZOTU1682 | | | | |
| | **Biofilm 1** | | **Biofilm 2** | | **Biofilm 3** | |
| | **Rel %** | **ZOTU** | **Rel %** | **ZOTU** | **Rel %** | **ZOTU** |
| | 0.01 | ZOTU49 | | | | |
| | 0.01 | ZOTU3965 | | | | |
| | 0.01 | ZOTU638 | | | | |
| | 0.01 | ZOTU4599 | | | | |
| T2 | 37.8 | ZOTU1024 | 58.0 | ZOTU1024 | 51.0 | ZOTU1024 |
| | 29.6 | ZOTU32 | 21.0 | ZOTU5147 | 23.0 | ZOTU11 |
| | 7.7 | ZOTU1581 | 11.2 | ZOTU5 | 10.5 | ZOTU5 |
| | 4.0 | ZOTU11 | 3.2 | ZOTU32 | 3.4 | ZOTU4295 |
| | 4.0 | ZOTU5 | 1.4 | ZOTU4295 | 3.2 | ZOTU6340 |
| | 3.9 | ZOTU2448 | 1.2 | ZOTU23 | 2.8 | ZOTU32 |
| | 3.6 | ZOTU145 | 1.0 | ZOTU274 | 1.5 | ZOTU103 |
| | 2.3 | ZOTU23 | 0.6 | ZOTU185 | 0.8 | ZOTU2448 |
| | 2.1 | ZOTU141 | 0.6 | ZOTU7 | 0.7 | ZOTU185 |
| | 2.0 | ZOTU4295 | 0.4 | ZOTU444 | 0.5 | ZOTU23 |
| | 1.7 | ZOTU185 | 0.2 | ZOTU141 | 0.4 | ZOTU2071 |
| | 0.3 | ZOTU7 | 0.2 | ZOTU148 | 0.4 | ZOTU43 |
| | 0.2 | ZOTU5147 | 0.2 | ZOTU2448 | 0.2 | ZOTU39 |
| | 0.2 | ZOTU444 | 0.1 | ZOTU42 | 0.2 | ZOTU7874 |
| | 0.1 | ZOTU39 | 0.1 | ZOTU2436 | 0.2 | ZOTU142 |
| | 0.09 | ZOTU7874 | 0.09 | ZOTU394 | 0.2 | ZOTU7 |
| | 0.09 | ZOTU421 | 0.09 | ZOTU113 | 0.2 | ZOTU290 |
| | 0.06 | ZOTU94 | 0.06 | ZOTU489 | 0.2 | ZOTU4868 |
| | 0.04 | ZOTU43 | 0.05 | ZOTU11702 | 0.1 | ZOTU2355 |
| | 0.04 | ZOTU148 | 0.04 | ZOTU13720 | 0.1 | ZOTU145 |
| | 0.04 | ZOTU13720 | 0.04 | ZOTU5438 | 0.1 | ZOTU94 |
| | 0.03 | ZOTU113 | 0.04 | ZOTU3965 | 0.08 | ZOTU13720 |
| | 0.02 | ZOTU2436 | 0.03 | ZOTU46 | 0.07 | ZOTU141 |
| | 0.02 | ZOTU42 | 0.03 | ZOTU7874 | 0.05 | ZOTU5147 |
| | 0.01 | ZOTU46 | 0.02 | ZOTU401 | 0.04 | ZOTU113 |
| | 0.01 | ZOTU2628 | 0.01 | ZOTU13630 | 0.04 | ZOTU3965 |
| | 0.01 | ZOTU6340 | 0.01 | ZOTU4937 | 0.03 | ZOTU148 |
| | 0.01 | ZOTU10313 | 0.01 | ZOTU10002 | 0.02 | ZOTU444 |
| | 0.01 | ZOTU142 | 0.01 | ZOTU4 | 0.02 | ZOTU42 |
| | 0.01 | ZOTU2881 | 0.01 | ZOTU5725 | 0.02 | ZOTU2436 |
| | 0.01 | ZOTU99 | | | 0.02 | ZOTU31 |
| | 0.01 | ZOTU1053 | | | 0.01 | ZOTU274 |
| | 0.01 | ZOTU125 | | | 0.01 | ZOTU5725 |
| | | | | | 0.01 | ZOTU10854 |
| | | | | | 0.01 | ZOTU10313 |
| | | | | | 0.01 | ZOTU74 |
| | | | | | 0.01 | ZOTU223 |
| | | | | | 0.01 | ZOTU951 |
| | | | | | 0.01 | ZOTU13071 |
| | | | | | 0.01 | ZOTU165 |
| | 90.3 | ZOTU4 | 86.5 | ZOTU4 | 34.5 | ZOTU46 |
| | 2.1 | ZOTU25 | 3.0 | ZOTU5147 | 21.5 | ZOTU1024 |
| | 2.0 | ZOTU28 | 2.9 | ZOTU25 | 14.9 | ZOTU4 |
| | 1.6 | ZOTU5147 | 2.0 | ZOTU1581 | 6.0 | ZOTU20 |
| | 1.1 | ZOTU1581 | 1.5 | ZOTU28 | 5.2 | ZOTU5147 |
| | 0.6 | ZOTU27 | 1.1 | ZOTU27 | 5.2 | ZOTU6049 |
| | 0.3 | ZOTU7 | 0.5 | ZOTU69 | 3.0 | ZOTU3965 |

| | | | | | | |
|---|---|---|---|---|---|---|
| | 0.3 | ZOTU11702 | 0.3 | ZOTU7 | 2.7 | ZOTU27 |
| | 0.3 | ZOTU69 | 0.3 | ZOTU11702 | 2.4 | ZOTU2448 |
| | 0.2 | ZOTU214 | 0.3 | ZOTU214 | 1.8 | ZOTU11702 |
| | **Biofilm 1** | | **Biofilm 2** | | **Biofilm 3** | |
| | **Rel %** | **ZOTU** | **Rel %** | **ZOTU** | **Rel %** | **ZOTU** |
| **T3** | 0.1 | ZOTU4295 | 0.2 | ZOTU3965 | 0.5 | ZOTU77 |
| | 0.1 | ZOTU23 | 0.2 | ZOTU20 | 0.4 | ZOTU69 |
| | 0.1 | ZOTU176 | 0.1 | ZOTU187 | 0.4 | ZOTU274 |
| | 0.1 | ZOTU187 | 0.1 | ZOTU176 | 0.3 | ZOTU7 |
| | 0.1 | ZOTU3965 | 0.1 | ZOTU4295 | 0.2 | ZOTU23 |
| | 0.09 | ZOTU13679 | 0.1 | ZOTU23 | 0.2 | ZOTU185 |
| | 0.08 | ZOTU20 | 0.1 | ZOTU13679 | 0.1 | ZOTU444 |
| | 0.05 | ZOTU5 | 0.08 | ZOTU2189 | 0.09 | ZOTU5 |
| | 0.05 | ZOTU148 | 0.08 | ZOTU4844 | 0.09 | ZOTU4295 |
| | 0.05 | ZOTU6049 | 0.08 | ZOTU6049 | 0.09 | ZOTU141 |
| | 0.04 | ZOTU1024 | 0.06 | ZOTU11380 | 0.08 | ZOTU11189 |
| | 0.04 | ZOTU11380 | 0.06 | ZOTU5 | 0.07 | ZOTU12758 |
| | 0.04 | ZOTU12603 | 0.06 | ZOTU1024 | 0.04 | ZOTU39 |
| | 0.03 | ZOTU113 | 0.05 | ZOTU274 | 0.04 | ZOTU155 |
| | 0.03 | ZOTU4844 | 0.04 | ZOTU42 | 0.04 | ZOTU113 |
| | 0.02 | ZOTU2189 | 0.04 | ZOTU141 | 0.03 | ZOTU1816 |
| | 0.02 | ZOTU49 | 0.03 | ZOTU148 | 0.03 | ZOTU148 |
| | 0.02 | ZOTU2436 | 0.03 | ZOTU12603 | 0.03 | ZOTU28 |
| | 0.02 | ZOTU274 | 0.03 | ZOTU809 | 0.03 | ZOTU5725 |
| | 0.01 | ZOTU207 | 0.02 | ZOTU46 | 0.03 | ZOTU4945 |
| | 0.01 | ZOTU42 | 0.02 | ZOTU5138 | 0.02 | ZOTU13679 |
| | 0.01 | ZOTU141 | 0.02 | ZOTU113 | 0.02 | ZOTU421 |
| | 0.01 | ZOTU155 | 0.02 | ZOTU2436 | 0.02 | ZOTU2436 |
| | 0.01 | ZOTU3774 | 0.02 | ZOTU155 | 0.02 | ZOTU42 |
| | 0.01 | ZOTU46 | 0.01 | ZOTU43 | 0.02 | ZOTU4844 |
| | 0.01 | ZOTU43 | 0.01 | ZOTU615 | 0.02 | ZOTU51 |
| | 0.01 | ZOTU11704 | 0.01 | ZOTU5438 | 0.01 | ZOTU43 |
| | 0.01 | ZOTU809 | 0.01 | ZOTU12758 | 0.01 | ZOTU144 |
| | 0.01 | ZOTU165 | 0.01 | ZOTU3774 | 0.01 | ZOTU165 |
| | 0.01 | ZOTU5138 | 0.01 | ZOTU110 | 0.01 | ZOTU74 |
| | | | 0.01 | ZOTU1682 | 0.01 | ZOTU1581 |
| | | | 0.01 | ZOTU32 | 0.01 | ZOTU11380 |
| | | | 0.01 | ZOTU49 | 0.01 | ZOTU6560 |
| | | | 0.01 | ZOTU1804 | 0.01 | ZOTU49 |
| | | | 0.01 | ZOTU4599 | 0.01 | ZOTU151 |

**Table S5** Shared taxa between biofilm communities and their initial original drinking water communities. Biofilms were grown on EPDM coupons in triplicate microcosm set ups. Original drinking water communities were different (B1, B2, T1, T2, T3). Colored rows highlight taxa that were present in both biofilm and original water.

| | | **Biofilm communities** | | | | | **Original drinking water communities** | | | | |
|---|---|---|---|---|---|---|---|---|---|---|---|
| | | 1 (Rel%) | 2 (Rel%) | 3 (Rel%) | Average (Rel%) | Deviation (Rel%) | 1 (Rel%) | 2 (Rel%) | 3 (Rel%) | Average (Rel%) | Deviation (Rel%) |
| **B1** | ZOTU5147 | 16.6 | 36.8 | 51.6 | 35.0 | 14.3 | 0.1 | 0.08 | 0.1 | 0.1 | 0.02 |
| | ZOTU10 | 49.6 | 0 | 12.4 | 20.7 | 21.1 | 0 | 0 | 0 | 0 | 0 |
| | ZOTU1024 | 0 | 42.2 | 0 | 14.1 | 19.9 | 0.2 | 0.01 | 0.16 | 0.1 | 0.08 |
| | ZOTU46 | 29.3 | 0 | 10.8 | 13.4 | 12.1 | 3.1 | 3.6 | 4.1 | 3.6 | 0.4 |
| | ZOTU35 | 0 | 6.4 | 14.2 | 6.9 | 5.8 | 0.01 | 0 | 0 | 0 | 0 |
| | ZOTU2255 | 0 | 4.2 | 8.1 | 4.1 | 3.3 | 0.05 | 0.01 | 0.05 | 0.04 | 0.02 |
| | ZOTU21 | 0 | 3.5 | 0 | 1.2 | 1.6 | 0 | 0 | 0 | 0 | 0 |
| | ZOTU61 | 1.6 | 0 | 0 | 0.5 | 0.7 | 0.01 | 0 | 0 | 0 | 0 |
| | ZOTU9849 | 0.8 | 0 | 0 | 0.3 | 0.4 | 30.9 | 31.7 | 26.7 | 29.8 | 2.2 |
| **B2** | ZOTU46 | 84.6 | 78.5 | 49.3 | 70.8 | 15.4 | 8.1 | 8.8 | 7.8 | 8.2 | 0.4 |
| | ZOTU5438 | 2.6 | 9.0 | 32.8 | 14.8 | 13.0 | 0.3 | 0.2 | 0.24 | 0.2 | 0.01 |
| | ZOTU21 | 5.5 | 5.7 | 10.2 | 7.1 | 2.2 | 0 | 0.01 | 0 | 0.01 | 0.01 |
| | ZOTU31 | 4.3 | 3.2 | 0 | 2.5 | 1.8 | 0 | 0 | 0 | 0 | 0 |
| | ZOTU5138 | 1.3 | 0 | 4 | 1.8 | 1.7 | 8.0 | 8.5 | 7.6 | 8.0 | 0.4 |
| | ZOTU77 | 0 | 1.8 | 1.5 | 1.1 | 0.8 | 0.05 | 0.08 | 0.09 | 0.07 | 0.02 |
| **T1** | ZOTU1024 | 56.7 | 76.9 | 76.8 | 70.2 | 9.5 | 0.03 | 0.03 | 0.04 | 0.03 | 0.01 |
| | ZOTU34 | 15.2 | 0 | 0 | 5.1 | 7.2 | 0 | 0 | 0 | 0 | 0 |
| | ZOTU9139 | 13.5 | 0 | 0 | 4.5 | 6.4 | 0.06 | 0.04 | 0.01 | 0.04 | 0.02 |
| | ZOTU4295 | 3.1 | 3.4 | 3.2 | 3.2 | 0.2 | 0.02 | 0.01 | 0 | 0.01 | 0.01 |
| | ZOTU25 | 0 | 5.0 | 4.3 | 3.1 | 2.2 | 0.01 | 0.04 | 0.03 | 0.02 | 0.01 |
| | ZOTU13 | 3.9 | 0 | 5.3 | 3.1 | 2.2 | 0 | 0 | 0.01 | 0 | 0 |
| | ZOTU94 | 0 | 4.0 | 2.6 | 2.2 | 1.7 | 0 | 0 | 0 | 0 | 0 |
| | ZOTU185 | 0 | 3.5 | 0 | 1.2 | 1.6 | 0 | 0 | 0.01 | 0 | 0 |
| **T2** | ZOTU1024 | 37.8 | 58.0 | 51.0 | 48.9 | 8.4 | 0.1 | 0.02 | 0.01 | 0.05 | 0.05 |
| | ZOTU32 | 29.6 | 3.2 | 0 | 10.9 | 13.3 | 0 | 0.01 | 0 | 0 | 0 |
| | ZOTU11 | 4.0 | 0 | 23.0 | 9.0 | 10.0 | 0.01 | 0 | 0 | 0 | 0 |
| | ZOTU5 | 4.0 | 11.2 | 10.5 | 8.6 | 3.2 | 0 | 0 | 0 | 0 | 0 |
| | ZOTU5147 | 0 | 21.0 | 0 | 7.0 | 9.9 | 0.06 | 0.00 | 0.01 | 0.02 | 0.02 |
| | ZOTU1581 | 7.7 | 0 | 0 | 2.6 | 3.6 | 0 | 0 | 0.01 | 0 | 0 |
| | ZOTU4295 | 0 | 1.4 | 3.4 | 1.6 | 1.4 | 0 | 0.01 | 0 | 0 | 0 |
| | ZOTU6340 | 0 | 0 | 3.2 | 1.1 | 1.5 | 0.02 | 0.01 | 0 | 0.01 | 0.01 |
| | ZOTU4 | 90.3 | 86.5 | 14.9 | 63.9 | 34.7 | 0 | 0.01 | 0 | 0 | 0 |
| | ZOTU46 | 0 | 0 | 34.5 | 11.5 | 16.2 | 0.02 | 0.01 | 0 | 0.01 | 0.01 |

| | | | | | | | | | | | |
|---|---|---|---|---|---|---|---|---|---|---|---|
| | ZOTU1024 | 0 | 0 | 21.5 | 7.2 | 10.1 | 0.2 | 0.01 | 0.01 | 0.07 | 0.08 |
| | ZOTU5147 | 1.6 | 3.0 | 5.2 | 3.3 | 1.5 | 0.03 | 0.00 | 0 | 0.01 | 0.01 |
| | | **Biofilm communities** | | | | | **Original drinking water communities** | | | | |
| | | 1 (Rel%) | 2 (Rel%) | 3 (Rel%) | Average (Rel%) | Deviation (Rel%) | 1 (Rel%) | 2 (Rel%) | 3 (Rel%) | Average (Rel%) | Deviation (Rel%) |
| **T3** | ZOTU20 | 0 | 0 | 6.0 | 2.0 | 2.8 | 0 | 0 | 0 | 0 | 0 |
| | ZOTU25 | 2.1 | 2.9 | 0 | 1.7 | 1.2 | 0 | 0 | 0 | 0 | 0 |
| | ZOTU28 | 2.0 | 1.5 | 0 | 1.2 | 0.8 | 0 | 0 | 0 | 0 | 0 |
| | ZOTU1581 | 1.1 | 2.0 | 0 | 1.0 | 0.8 | 0 | 0 | 0.01 | 0 | 0 |

**Table S6** Relative abundances of taxonomically assigned families within biofilm communities that formed on EPDM coupons and with different original starting communities (B1, B2, T1, T2, T3).

| | **Biofilm 1** | | **Biofilm 2** | | **Biofilm 3** | |
|---|---|---|---|---|---|---|
| | **Rel %** | **Family** | **Rel %** | **Family** | **Rel %** | **Family** |
| **B1** | 49.6 | Xanthomonadaceae | 80.9 | Comamonadaceae | 52.4 | Comamonadaceae |
| | 29.3 | Burkholderiaceae | 6.4 | Rhizobiaceae | 14.2 | Rhizobiaceae |
| | 18.2 | Comamonadaceae | 4.2 | Moraxellaceae | 12.4 | Xanthomonadaceae |
| | 1.6 | Phyllobacteriaceae | 3.5 | Hyphomonadaceae | 10.8 | Burkholderiaceae |
| | 0.6 | NA | 2.2 | Xanthomonadaceae | 8.1 | Moraxellaceae |
| | 0.2 | Uncultured_bacterium | 1.4 | Nocardioidaceae | 0.8 | NA |
| | 0.2 | Methylobacteriaceae | 0.6 | Caulobacteraceae | 0.7 | Phyllobacteriaceae |
| | 0.1 | Bradyrhizobiaceae | 0.3 | Burkholderiaceae | 0.3 | Methylobacteriaceae |
| | 0.04 | Bacillaceae | 0.2 | Methylobacteriaceae | 0.2 | Nocardioidaceae |
| | 0.03 | Caulobacteraceae | 0.1 | Bradyrhizobiaceae | 0.2 | Bradyrhizobiaceae |
| | 0.03 | Brucellaceae | 0.07 | NA | 0.04 | Bacillaceae |
| | 0.02 | Rhizobiaceae | 0.07 | Phyllobacteriaceae | 0.01 | Brucellaceae |
| | 0.02 | Sphingomonadaceae | 0.02 | Brucellaceae | 0.01 | Sphingomonadaceae |
| | 0.01 | Hyphomonadaceae | 0.02 | Bacillaceae | 0.01 | Hyphomonadaceae |
| | | | 0.01 | Sphingomonadaceae | | |
| **B2** | 84.6 | Burkholderiaceae | 78.5 | Burkholderiaceae | 49.3 | Burkholderiaceae |
| | 5.6 | Hyphomonadaceae | 10.5 | Comamonadaceae | 38.0 | Comamonadaceae |
| | 4.3 | Caulobacteraceae | 5.7 | Hyphomonadaceae | 10.2 | Hyphomonadaceae |
| | 4.3 | Comamonadaceae | 3.2 | Caulobacteraceae | 1.6 | NA |
| | 0.4 | Methylobacteriaceae | 1.8 | NA | 0.6 | Caulobacteraceae |
| | 0.3 | NA | 0.1 | Methylobacteriaceae | 0.2 | Methylobacteriaceae |
| | 0.2 | Cytophagaceae | 0.05 | Bradyrhizobiaceae | 0.09 | Bradyrhizobiaceae |
| | 0.1 | Bradyrhizobiaceae | 0.02 | Sphingomonadaceae | 0.02 | Bacillaceae |
| | 0.05 | Sphingomonadaceae | 0.01 | Cytophagaceae | 0.02 | Brucellaceae |
| | 0.03 | Bacillaceae | | | 0.01 | Xanthomonadaceae |
| | 0.02 | Brucellaceae | | | 0.01 | Sphingomonadaceae |
| | 0.01 | Rhodocyclaceae | | | 0.01 | Chitinophagaceae |
| | 0.01 | Oxalobacteraceae | | | 0.01 | Flavobacteriaceae |
| | 0.01 | Rhizobiaceae | | | | |
| | 0.01 | Microbacteriaceae | | | | |
| **T1** | 56.9 | Comamonadaceae | 78.4 | Comamonadaceae | 76.9 | Comamonadaceae |
| | 15.3 | Xanthomonadaceae | 5.0 | Caulobacteraceae | 5.3 | Chitinophagaceae |
| | 13.5 | Leptospiraceae | 4.0 | Xanthomonadaceae | 4.3 | Caulobacteraceae |
| | 3.9 | Chitinophagaceae | 3.5 | Rhodocyclaceae | 3.2 | Rhodocyclaceae |
| | 3.1 | Rhodocyclaceae | 3.5 | Rhodobacteraceae | 3.1 | Rhodobacteraceae |
| | 1.8 | Rhodobacteraceae | 2.5 | Chitinophagaceae | 2.6 | Xanthomonadaceae |
| | 1.5 | Rhodospirillales_Incer-tae_Sedis | 1.2 | NA | 1.3 | NA |
| | 1.1 | Bradyrhizobiaceae | 1.0 | Bradyrhizobiaceae | 1.1 | Sphingomonadaceae |
| | 0.9 | NA | 0.2 | Rhodospirrales_Incer-tae_Sedis | 1.0 | Rhodospirrales_Incertae_Sedis |
| | 0.7 | Caulobacteraceae | 0.2 | Sphingomonadaceae | 0.9 | Bradyrhizobiaceae |
| | 0.5 | Sphingomonadaceae | 0.2 | Methylobacteriaceae | 0.2 | Methylobacteriaceae |
| | 0.4 | Methylobacteriaceae | 0.1 | Microbacteriaceae | 0.04 | Microbacteriaceae |

| | | | | | | |
|---|---|---|---|---|---|---|
| | 0.2 | Microbacteriaceae | 0.1 | Nitrosomonadaceae | 0.03 | Brucellaceae |
| | 0.1 | Xanthobacteraceae | 0.04 | uncultured_bacterium | 0.03 | Nitrosomonadaceae |
| | **Biofilm 1** | | **Biofilm 2** | | **Biofilm 3** | |
| | **Rel %** | **Family** | **Rel %** | **Family** | **Rel %** | **Family** |
| | 0.07 | Bacillaceae | 0.03 | Thermaceae | 0.03 | Bacillaceae |
| | 0.07 | Nitrosomonadaceae | 0.03 | Xanthobacteraceae | 0.02 | uncultured_bacterium |
| | 0.07 | uncultured_bacterium | 0.01 | Propionibacteriaceae | 0.02 | Xanthobacteraceae |
| | 0.02 | Thermaceae | | | 0.01 | Burkholderiaceae |
| | 0.02 | Brucellaceae | | | 0.01 | Leptospiraceae |
| | 0.01 | Burkholderiaceae | | | 0.01 | Thermaceae |
| | 0.01 | Anaerolineaceae | | | | |
| | 0.01 | Nanoarchaeota_archaeon_SCGC_AAA011-D5 | | | | |
| **T2** | 48.3 | Comamonadaceae | 79.8 | Comamonadaceae | 54.6 | Comamonadaceae |
| | 29.6 | Sphingomonadaceae | 11.2 | Chitinophagaceae | 23.0 | Caulobacteraceae |
| | 4.0 | Caulobacteraceae | 3.2 | Sphingomonadaceae | 10.5 | Chitinophagaceae |
| | 4.0 | Chitinophagaceae | 1.4 | Rhodocyclaceae | 3.4 | Rhodocyclaceae |
| | 3.9 | Rhodospirillales_Incer-tae_Sedis | 1.2 | Bradyrhizobiaceae | 2.8 | Sphingomonadaceae |
| | 3.7 | Xanthomonadaceae | 1.0 | Rhizobiales_Incer-tae_Sedis | 1.8 | NA |
| | 2.3 | Bradyrhizobiaceae | 0.6 | Rhodobacteraceae | 1.1 | Rhodospirillales_Incertae_Sedis |
| | 2.0 | Rhodocyclaceae | 0.6 | Methylobacteriaceae | 0.7 | Bradyrhizobiaceae |
| | 1.7 | Rhodobacteraceae | 0.2 | Bacillaceae | 0.7 | Rhodobacteraceae |
| | 0.3 | Methylobacteriaceae | 0.2 | Rhodospirillales_Incer-tae_Sedis | 0.4 | TK34 |
| | 0.1 | NA | 0.1 | NA | 0.2 | Xanthomonadaceae |
| | 0.05 | Bacillaceae | 0.1 | I-10 | 0.2 | Methylobacteriaceae |
| | 0.02 | Brucellaceae | 0.1 | Brucellaceae | 0.2 | MNG7 |
| | 0.01 | Burkholderiaceae | 0.06 | Rhodospirillaceae | 0.2 | Legionellaceae |
| | 0.01 | Propionibacteriaceae | 0.03 | Burkholderiaceae | 0.1 | Parviterribacteraceae |
| | 0.01 | uncultured | 0.02 | Oxalobacteraceae | 0.03 | Bacillaceae |
| | 0.01 | Xanthobacteraceae | 0.01 | uncultured_bacterium | 0.02 | Brucellaceae |
| | | | | | 0.01 | Rhizobiales_Incer-tae_Sedis |
| | | | | | 0.01 | Microbacteriaceae |
| | | | | | 0.01 | uncultured_bacterium |
| **T3** | 90.4 | NA | 86.8 | NA | 34.5 | Burkholderiaceae |
| | 3.2 | Comamonadaceae | 5.6 | Comamonadaceae | 29.0 | Comamonadaceae |
| | 2.1 | Caulobacteraceae | 3.0 | Caulobacteraceae | 18.4 | NA |
| | 2.0 | Flavobacteriaceae | 1.5 | Flavobacteriaceae | 6.0 | Xanthomonadaceae |
| | 0.6 | Sphingomonadaceae | 1.1 | Sphingomonadaceae | 5.2 | Caulobacteraceae |
| | 0.3 | Methylobacteriaceae | 0.5 | ODP1230B8.23 | 2.7 | Sphingomonadaceae |
| | 0.3 | ODP1230B8.23 | 0.3 | Methylobacteriaceae | 2.4 | Rhodospirillales_Incertae_Sedis |
| | 0.2 | env.OPS_17 | 0.3 | env.OPS_17 | 0.4 | ODP1230B8.23 |
| | 0.2 | I-10 | 0.3 | I-10 | 0.4 | Rhizobiales_Incer-tae_Sedis |
| | 0.1 | Rhodocyclaceae | 0.2 | Xanthomonadaceae | 0.3 | Methylobacteriaceae |
| | 0.1 | Bradyrhizobiaceae | 0.1 | Rhodocyclaceae | 0.3 | Rhodobacteraceae |

| | | | | | | |
|---|---|---|---|---|---|---|
| | 0.08 | Xanthomonadaceae | 0.1 | Bradyrhizobiaceae | 0.2 | Bradyrhizobiaceae |
| | 0.05 | Chitinophagaceae | 0.08 | Phycisphaeraceae | 0.09 | Chitinophagaceae |
| | 0.05 | Bacillaceae | 0.06 | Chitinophagaceae | 0.09 | Rhodocyclaceae |
| | **Biofilm 1** | | **Biofilm 2** | | **Biofilm 3** | |
| | **Rel %** | **Family** | **Rel %** | **Family** | **Rel%** | **Family** |
| | 0.03 | Opitutaceae | 0.05 | Rhizobiales_Incer-tae_Sedis | 0.04 | Hyphomicrobiaceae |
| | 0.02 | Phycisphaeraceae | 0.04 | Bacillaceae | 0.03 | Bacillaceae |
| | 0.02 | Brucellaceae | 0.04 | Opitutaceae | 0.03 | Flavobacteriaceae |
| | 0.02 | Rhizobiales_Incer-tae_Sedis | 0.02 | Burkholderiaceae | 0.02 | Brucellaceae |
| | 0.01 | Burkholderiaceae | 0.02 | Brucellaceae | 0.02 | Microbacteriaceae |
| | 0.01 | Hyphomicrobiaceae | 0.02 | Hyphomicrobiaceae | 0.01 | uncultured_bacteriu m |
| | 0.01 | Microbacteriaceae | 0.01 | Rhodospirillales_Incer-tae_Sedis | | |
| | 0.01 | Rhodospirillales_Incer-tae_Sedis | 0.01 | Rhodobacteraceae | | |
| | | | 0.01 | Rhodospirillaceae | | |

**Table S7** Attachment performances for selected biofilm and original drinking water communities. B1 (bottled water), B2 (bottled water), T1 (tapped groundwater), T2 (tap water), T3 (tap water). Experiments on EPDM were performed in experimental triplicates, glass controls were done in singletons.

| **Maximum attachment rate (TCC/cm$^2$/h)** | | | | |
|---|---|---|---|---|
| | EPDM | | Glass | |
| | **Original** | **Selected** | **Original** | **Selected** |
| B1 | 3.3±0.6 x $10^4$ (n = 3) | 5.4±0.1 x $10^4$ (n = 3) | 2.8 x $10^4$ | 5.3 x $10^4$ |
| B2 | 3.3±0.4 x $10^4$ (n = 3) | 5.0±0.7 x $10^4$ (n = 3) | 3.2 x $10^4$ | 4.3 x $10^4$ |
| T1 | 3.1±0.3 x $10^4$ (n = 3) | 7.8±0.5 x $10^4$ (n = 3) | 3.0 x $10^4$ | 6.5 x $10^4$ |
| T2 | 1.7±0.3 x $10^4$ (n = 3) | 2.9±0.1 x $10^4$ (n = 3) | 1.6 x $10^4$ | 2.3 x $10^4$ |
| T3 | 4.6±0.8 x $10^3$ (n = 3) | 3.9±0.4 x $10^4$ (n = 3) | 3.4 x $10^3$ | 3.1 x $10^4$ |
| | **Maximum attachment (TCC/cm$^2$)** | | | |
| | EPDM | | Glass | |
| | **Original** | **Selected** | **Original** | **Selected** |
| B1 | 6.3±0.1 x $10^4$ (n = 3) (80±1%) | 8.1±0.2 x $10^4$ (n = 3) (97±0%) | 1.4x $10^4$ (81%) | 4.8 x $10^3$ (92%) |
| B2 | 7.1±0.2 x $10^4$ (n = 3) (76±4%) | 6.5±0.1 x $10^4$ (n = 3) (83±2%) | 3.0 x $10^4$ (66%) | 1.1 x $10^4$ (87%) |
| T1 | 7.4±0.3 x $10^4$ (n = 3) (45±2%) | 1.2±0.0 x $10^5$ (n = 3) (87±1%) | 9.6 x $10^4$ (41%) | 3.2 x $10^4$ (76%) |
| T2 | 4.0±0.1 x $10^4$ (n = 3) (75±3%) | 5.0±0.0 x $10^4$ (n = 3) (92±1%) | 1.8 x $10^4$ (67%) | 7.7 x $10^3$ (86%) |
| T3 | 1.4±0.3 x $10^4$ (n = 3) (24±4%) | 6.7±0.1 x $10^4$ (n = 3) (94±1%) | 5.3 x $10^4$ (12%) | 5.4 x $10^3$ (92%) |

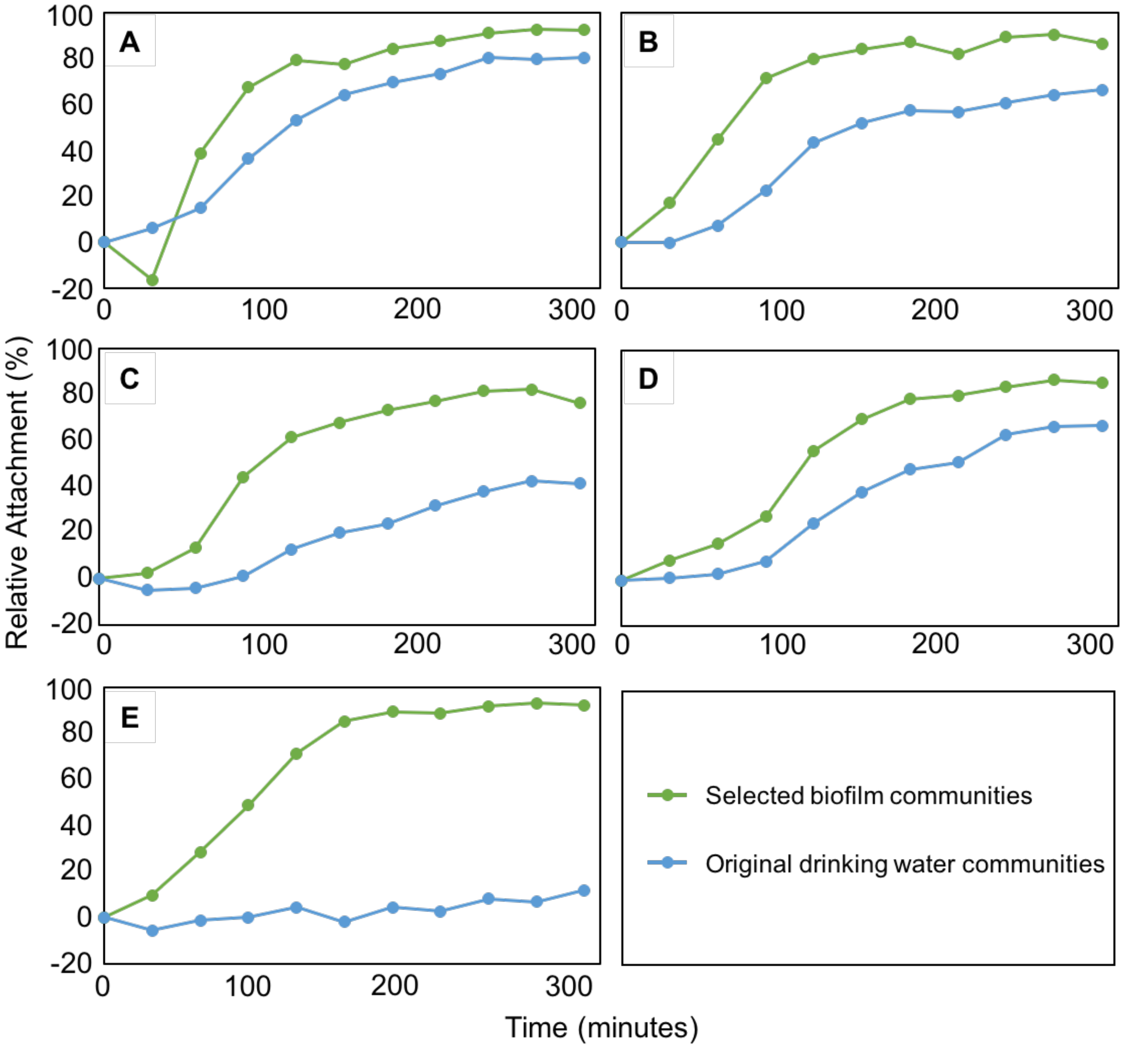

**Figure S1** Attachment performances of selected biofilm and original drinking water communities (initial colonization) on glass coupons. (A) bottled water *B1*, (B) bottled water *B2*, (C) tapped groundwater *T1*, (D) tap water *T2*, and (E) tap water *T3*. Starting concentrations between selected and original communities were adapted to be similar within set ups.
